# The rapid developmental rise of somatic inhibition disengages hippocampal dynamics from self-motion

**DOI:** 10.1101/2021.06.08.447542

**Authors:** Robin F. Dard, Erwan Leprince, Julien Denis, Shrisha Rao-Balappa, Dmitrii Suchkov, Richard Boyce, Catherine Lopez, Marie Giorgi-Kurz, Tom Szwagier, Theo Dumont, Hervé Rouault, Marat Minlebaev, Agnes Baude, Rosa Cossart, Michel A. Picardo

## Abstract

Early electrophysiological brain oscillations recorded in preterm babies and newborn rodents are initially mostly ignited by bottom-up sensorimotor activity and only later can detach from external inputs. This is a hallmark of most developing brain areas including the hippocampus, which in the adult brain, functions in integrating external inputs onto internal dynamics. Such developmental disengagement from external inputs is likely a fundamental step for the proper development of cognitive internal models. Despite its importance, the developmental timeline and circuit basis for this disengagement remain unknown. To address this issue, we have investigated the daily evolution of CA1 dynamics and underlying circuits during the first two postnatal weeks of mouse development using two-photon calcium imaging in non-anesthetized pups. We show that the first postnatal week ends with an abrupt shift in the representation of self-motion in CA1. Indeed, most CA1 pyramidal cells switch from activated to inhibited by self-generated movements at the end of the first postnatal week whereas the majority of GABAergic neurons remain positively modulated throughout this period. This rapid switch occurs within two days and follows the rapid anatomical and functional surge of local somatic GABAergic innervation. The observed change in dynamics is consistent with a two-population model undergoing a strengthening of inhibition. We propose that this abrupt developmental transition inaugurates the emergence of internal cognitive models.

## INTRODUCTION

The adult hippocampus serves multiple cognitive functions including navigation and memory. These functions rely on the ability of hippocampal circuits to integrate external inputs conveying multi-sensory, proprioceptive, contextual and emotional information onto internally-generated dynamics. Therefore, the capacity to produce internally coordinated neuronal activity detached from environmental inputs is central to the cognitive functions of the hippocampus such as planning and memory (Buzsáki, 2015; Buzsáki and Moser, 2013). In contrast to the adult situation, the developing hippocampus, like many developing cortical structures, is mainly driven by bottom-up external environmental and body-derived signals, including motor twitches generated in the spinal cord and/or the brainstem (Dooley et al., 2020; Inácio et al., 2016; Karlsson et al., 2006; Mohns and Blumberg, 2010; Rio-Bermudez et al., 2020; Valeeva et al., 2019a). These produce early sharp waves conveyed by inputs from the entorhinal cortex (Valeeva et al., 2019a). The emergence of self-organized sequences without reliance on external cues in the form of sharp wave ripples (SWRs) is only observed after the end of the second postnatal week and sequential reactivations even a week later (Farooq and Dragoi, 2019; Muessig et al., 2019). Therefore, early hippocampal activity as measured with electrophysiological recordings is first externally-driven while the emergence of internal dynamics is protracted. The timing and the circuit mechanisms of the switch between motion-guided and internally produced hippocampal dynamics remain unknown. They have been proposed to rely on the maturation of CA3 and extrinsic hippocampal inputs, however, a possible role of local connectivity, in particular recurrent somatic inhibition cannot be excluded (Cossart and Khazipov, 2021a).

Local GABAergic interneurons could be critically involved in this phenomenon for several reasons. First, both theoretical and experimental work suggest that self-organized internal neuronal network dynamics require feedback connections to produce an emergent state of activity independently from the incoming input (Hopfield and Tank, 2005; Hopfield, 1982; Yuste, 2015). Feedback circuits are mainly GABAergic in CA1 (but not necessarily inhibitory), given the scarcity of recurrent glutamatergic connections in that hippocampal sub-region (Bezaire and Soltesz, 2013). Second, GABAergic interneurons, in particular the perisomatic subtypes, are long known to shape the spatial and temporal organization of internal CA1 dynamics (Buzsáki, 2015; Lee et al., 2014; Soltesz and Losonczy, 2018; Valero et al., 2015). However, GABAergic perisomatic cells display a delayed maturation profile both at structural (Jiang et al., 2001; Marty et al., 2002; Morozov and Freund, 2003; Tyzio et al., 1999) and functional levels (Ben-Ari, 2002; Doischer et al., 2008; Jiang et al., 2001; Khazipov et al., 2004; Marty et al., 2002; Morozov and Freund, 2003; Murata and Colonnese, 2020; Tyzio et al., 1999), and the precise developmental timeline for their postnatal development remains unknown, partly due to the difficulty in labeling them (Donato et al., 2017).

Here we investigate the evolution of CA1 dynamics during the first and second postnatal weeks of mouse development with an eye on the specific patterning of activity of CA1 GABAergic neurons. To this aim, we adapted two-photon calcium imaging of CA1 dynamics using virally-expressed GCaMP6 through a cranial window in non-anesthetized pups. We show that the first postnatal week ends with an abrupt switch in the representation of self-motion in CA1: principal neurons were synchronized by spontaneous movement before P9, whereas self-motion decreased their activity after that time point. Consistent with a two-population neuronal model, this switch was locally paralleled by the rapid anatomical and functional surge of somatic GABAergic interneurons and no significant change in external inputs. Self-generated bottom-up inputs may thus directly contribute to the emergence of somatic GABAergic inhibition and in this way calibrate local circuits to the magnitude of external inputs prior to the opening of experience-dependent plasticity.

## RESULTS

### Progressive evolution of CA1 neuronal dynamics

In order to induce stable and early expression of the calcium indicator protein GCaMP6s, pups were injected with the AAV1-hSyn-GCaMP6s.WPRE.SV40 virus in the brain lateral ventricle on the day of birth (P0, **Figure 1A and Figure 1 - figure supplement 1A**). Five to twelve days after injection, the hippocampal CA1 region of non-anesthetized pups was imaged through a window implant placed on the same day (**Figure 1A, see methods**). This acute window implant did not alter hippocampal dynamics as revealed by *in vivo* bilateral silicon probes electrophysiological recordings of early sharp waves (eSW, **Figure 1 - figure supplement 1B**) in P6-8 and P11 pups (N=4 and N=2 respectively) expressing GCaMP6s (with a frequency of 2.6 eSW/min (25 % 1.15 eSW/min and 75 % 4.16 eSW/min) for the ipsilateral side and 3.49 eSW/min (25 % 1.96 eSW/min and 75 % 5.1 eSW/min, *p-value* = 0.39) for the contralateral side (**Figure 1 - figure supplement 1B**). eSW synchronization between hemispheres was also preserved (peak at 0.087 +/- 0.027, **Figure 1 - figure supplement 1B**) which is in agreement with previous data (Graf et al., 2021; Valeeva et al., 2019b). Finally, we checked for the presence of other types of oscillations in both hemispheres and observed a peak in the theta range in the P11 mouse pups in both hemispheres (Ipsi: peak at 4.3 Hz of amplitude 3.6×10^−3^ +/- 1.2×10^−3^ mV^2^/Hz; Contra: peak at 4.1 Hz of amplitude 2×10^−3^ +/- 3×10^−4^ mV^2^/Hz (jackknife std)). Therefore, the presence of the window implant preserved the previously described network activity of the early postnatal period in developing rodents. Thus, we pursued the description of early multineuron CA1 dynamics using calcium imaging (62 imaging sessions, 35 mouse pups aged between 5 and 12 days, yielding a total of 33,412 cells, see **Table 1** for details on each session and their inclusion in the figures).

**Figure 1:**
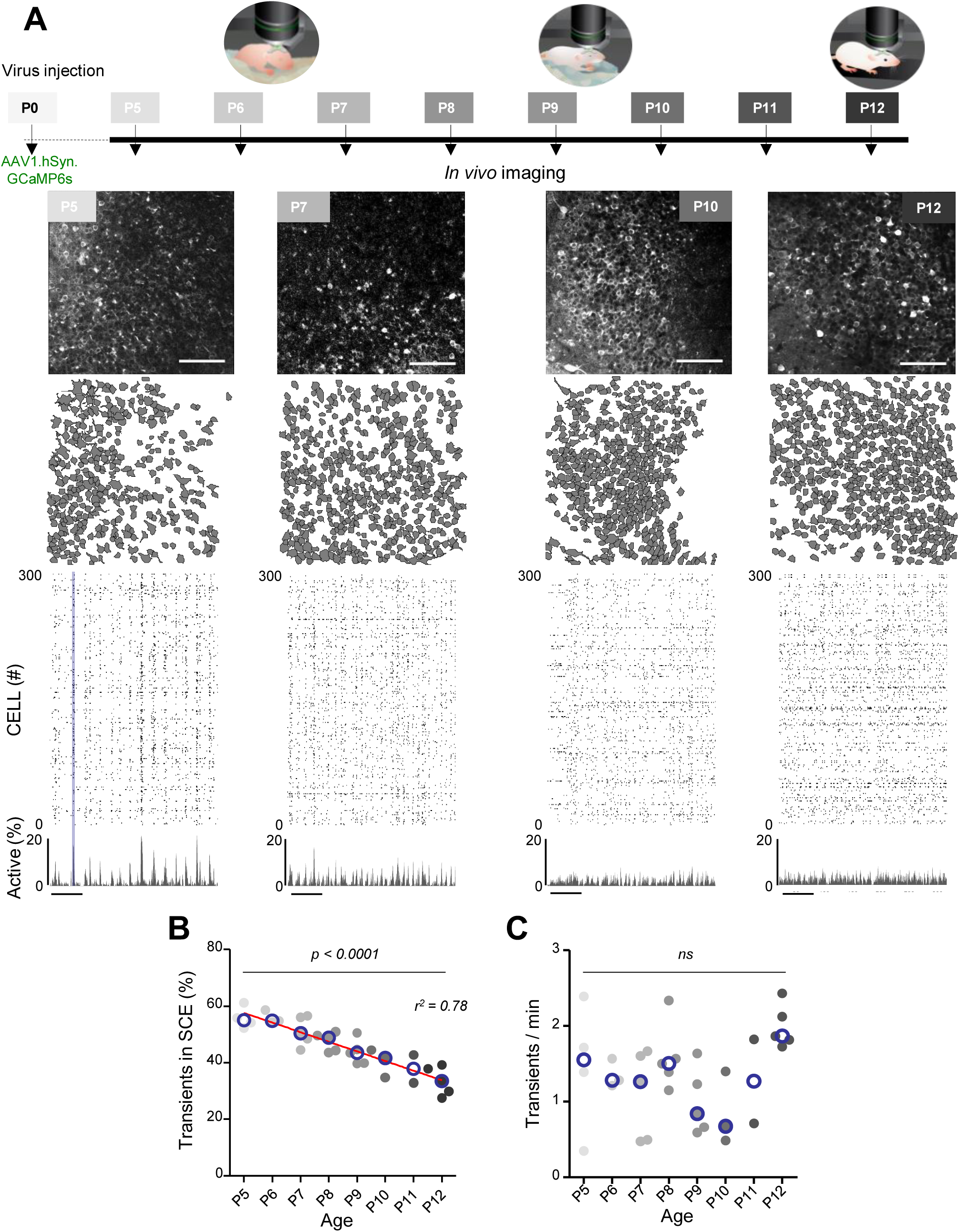
Evolution of CA1 dynamics during the first two postnatal weeks. **(A)**. Schematic of the experimental timeline. On postnatal day 0 (P0), 2μL of a non-diluted viral solution were injected in the left lateral ventricle of mouse pups. From five to twelve days after injection (P5-P12), acute surgery for window implantation above the corpus callosum was performed and followed by 2-photon calcium imaging recordings. Top panel: Four example recordings are shown to illustrate the imaging fields of view in the stratum pyramidale of the CA1 region of the hippocampus (scale bar: 100 μm). Middle panel: Contour maps showing the cells detected using Suite2p in the corresponding fields of view. Bottom panel: Raster plots inferred by DeepCINAC activity classifier, showing 300 randomly selected cells over the first 5 minutes of recording obtained for these imaging sessions (P5, P7, P10, P12, see full raster plots for these imaging sessions in Figure 1 - figure supplement 1D). In the raster plot from the P5 mouse the blue rectangle illustrates one synchronous calcium event (SCE). Scale bar for time is 1 min. CICADA configuration files to reproduce example rater plots and cell contours are available in Figure 1 - Source Data 1. **(B)**. Evolution of the ratio of calcium transients within SCEs over the total number of transients across age. Each dot represents a mouse pup and is color coded from light gray (P5) to black (P12), the open blue circles represent the median of the age group. The red line represents the linear fit of the data with r^2^=0.78 and p<0.0001. Results to build the distribution as well as CICADA configuration file to reproduce the analysis are available in Figure 1 - Source Data 1. **(C)**. Evolution of the number of transients per minute across the first two postnatal weeks. Each dot represents the mean transient frequency from all cells imaged in one animal and is color coded from light gray (P5) to black (P12). The open blue circles represent the median of the age group. No significant influence of the age was found (Kruskal-Wallis test, 8 groups, KW stat =12.63, p-value = 0.0816). Results to build the distribution as well as CICADA configuration file to reproduce the analysis are available in Figure 1 - Source Data 1.

**Table 1:**
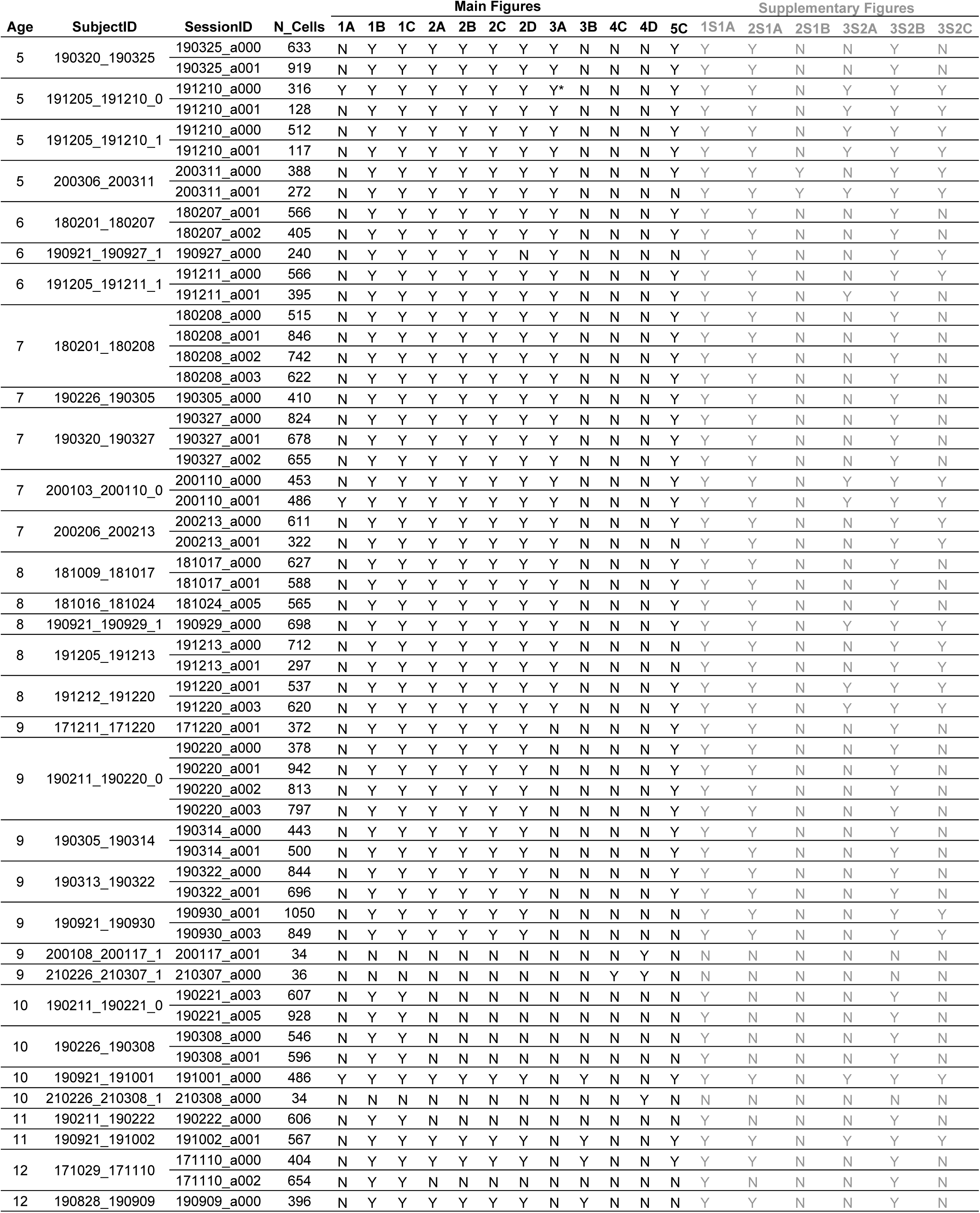
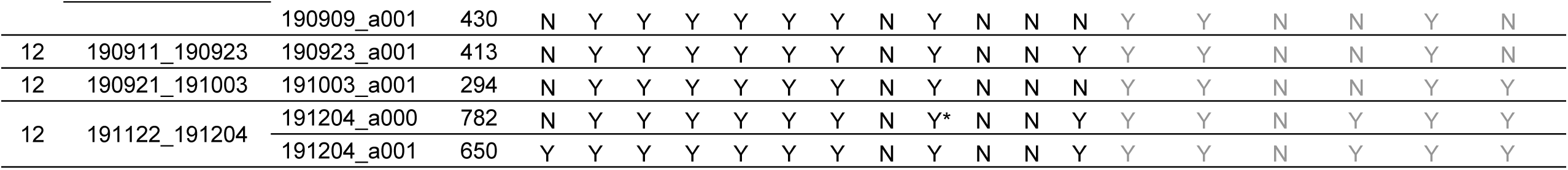
Details on the 62 imaging sessions from the 35 mouse pups showing the number of cells recorded in each session and how they were used in the figures. In main figure 3A, * shows mouse pups that are used for illustration. Y: included in the panel, N: not included.

The contours of the imaged neurons and their calcium fluorescence events were extracted using Suite2P (Pachitariu et al., 2017) and DeepCINAC (Denis et al., 2020), respectively. Representative examples of fields of view, contour maps and activity raster plots from recordings in P5, P7, P10 and P12 mouse pups are shown in Figure 1A. Consistent with previous electrophysiological studies (Leinekugel et al., 2002; Mohns and Blumberg, 2008; Valeeva et al., 2019a), spontaneous neuronal activity in the CA1 region of P5-6 pups alternated between recurring population bursts (Synchronous Calcium Events, SCEs), and periods of low activity (**Figure 1A**). In P5-6 mouse pups more than half of the detected calcium transients occurred within SCEs (P5: median value 55 % N=4, n=8; P6: median value 55 % N=3, n=5; N: mice, n: imaging sessions, **Figure 1B**). Activity then became progressively continuous as evidenced by the linear decrease in the proportion of calcium transients occurring during SCEs (r^2^=0.78, p<0.0001) to finally reach 33 % in P12 mouse pups (P12: N=5, n=8, **Figure 1B**). These changes in CA1 dynamics cannot be explained by a change in the activation rate of individual neurons as this parameter did not significatively change during the developmental period studied here (**Figure 1C**). Even though there was globally no significant change in the rate of neuronal activation as measured with calcium imaging from P5 to P12, we observed a small, but not significant decrease at P9-10 consistent with the description of a transient period of ‘neural quiescence’ at the beginning of the second postnatal week (Dominguez et al., 2021). We conclude that CA1 dynamics progressively evolve from discontinuous to continuous during the first two postnatal weeks, in agreement with previous electrophysiological studies (Cossart and Khazipov, 2021b; Mohns and Blumberg, 2008; Valeeva et al., 2019a).

### Early synchronous calcium events correlate with spontaneous motor activity

Previous extracellular electrophysiological recordings indicated that, in developing rodents, CA1 dynamics followed spontaneous motor activity during the first postnatal week (Karlsson et al., 2006; Rio-Bermudez et al., 2020; Valeeva et al., 2019a). Hence, we next examined the relationship between population activity and movement as monitored using either piezo recordings or infrared cameras (see methods). We first computed peri-movement time histograms (PMTHs) plotting the fraction of active neurons centered on the onset of all movements. In mouse pups younger than P9, movements were followed by a significant increase in the percentage of active cells (**Figure 2A**, P5-8 median above chance level, **Supplementary movies 1-3**). In contrast, after P9, movements were followed by a significant decrease of activity (**Figure 2A**, P10-12 median below chance level, **Supplementary Movies 4-6**). Short myoclonic movements such as twitches/startles, happening during periods of active sleep (Gramsbergen et al., 1970; Jouvet-Mounier and Astic, 1968; Karlsson et al., 2006) as opposed to longer movements, happening mostly during wakefulness, may induce different activity patterns in the hippocampus (Mohns and Blumberg, 2008). However, we found that both movement types did not significatively differ in their impact on CA1 activity (**Figure 2 - figure supplement 1A**). Furthermore calcium imaging was combined with nuchal EMG recordings in one P5 mouse pup, and an increase in the percentage of active cells was observed following movements occurring both during REM sleep and wakefulness (**Figure 2 - figure supplement 1B**). All movement types were thus combined in further analysis. Post-movement activity was next computed, as defined by the number of active cells in the two seconds following movement onset divided by the number of active cells within a four seconds-long time window centered on movement onset (see methods, **Figure 2B and Figure 2 - figure supplement 1C**). The median post-movement activity progressively decreased from P5 to P9 until it suddenly dropped at P10 and stabilized until P12. P9 marked the transition in the relationship between movement and CA1 activity. Indeed, the median post-movement activity exceeded 50 % from P5 to P8 (P5: 71 %, P6: 65 %, P7: 60 %, P8: 56 %). This is consistent with the evolution of PMTHs (**Figure 2A**). After P9, the median post-movement activity was lower than 50 % (P10: 39 %, P11: 35 %, P12: 40 %), thus revealing the inhibitory action of movement on activity. We next defined as ‘inhibiting movements’ all the movements with a post-movement activity lower than 40 % and computed their proportion in each mouse (**Figure 2C**). The proportion of ‘inhibiting’ movements was stable before P9 (P5: 11 %, P6: 16 %, P7: 10 %, P8: 15 %). Again, P9 marks a transition since we observed that approximately half of the movements were followed by an inhibition of CA1 activity in P10-12 mice (P10: 55 %, P11: 58 %, P12: 48 %). The proportion of ‘inhibiting’ movements varies with age as a sigmoïd function with P9 being the transition time point (V50=9.015, r^2^=0.75). In line with the emergence of movement-induced inhibition, the fraction of neurons significatively associated with immobility also increased with age (**Figure 2D**) suggesting an “internalization” of neuronal activity. Altogether, these results indicate that the end of the first postnatal week marks a transition in the evolution of CA1 dynamics, with both a decorrelation and an internalization of neuronal activity. We next investigated the circuit mechanisms supporting these changes.

**Figure 2:**
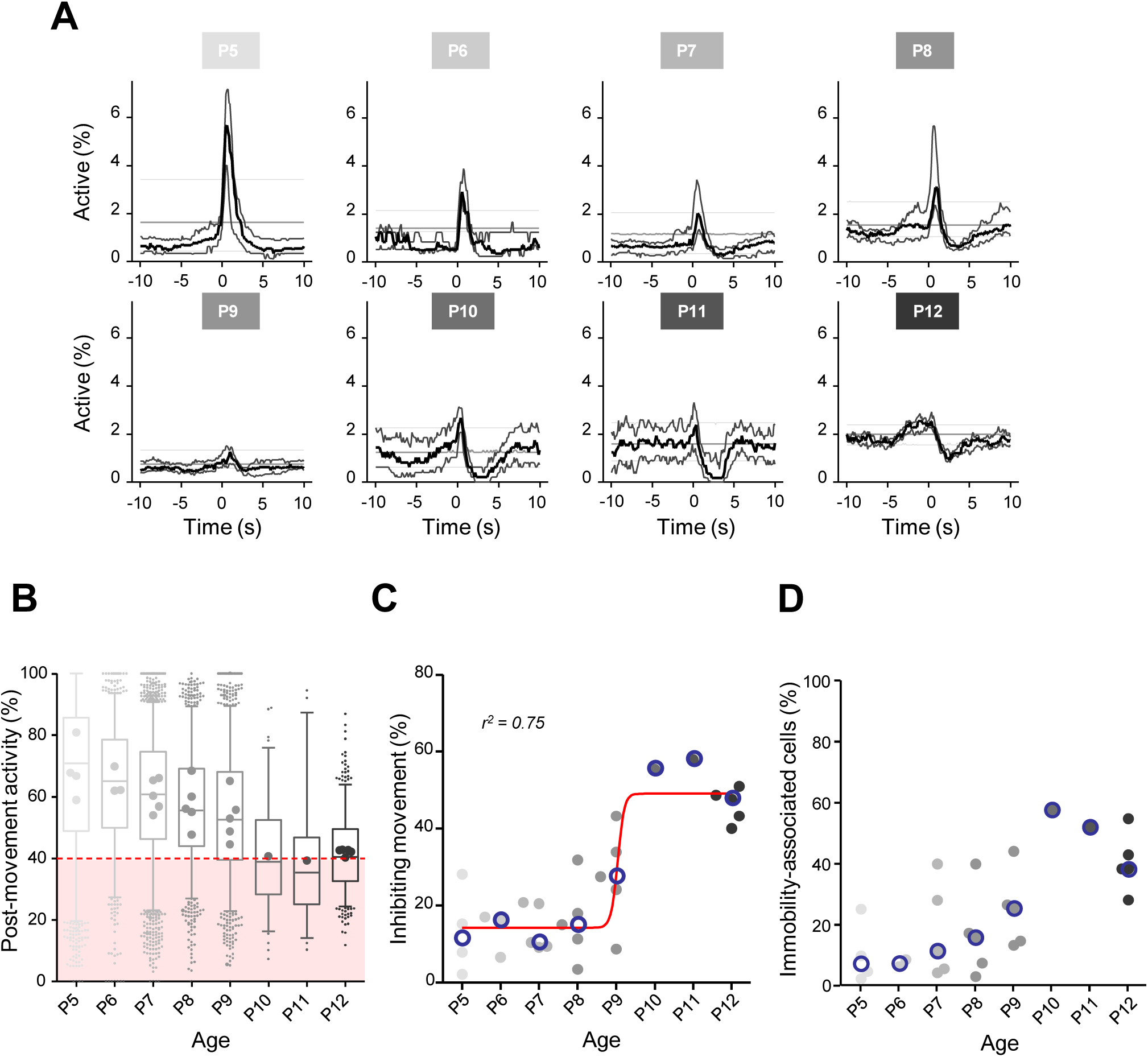
Linking CA1 dynamics to movement during the first two postnatal weeks. **(A)**. Peri-Movement-Time-Histograms (PMTH) representing the percentage of active cells centered on the onset of the mouse movements. The dark line indicates the median value, the two thick grey lines represent the 25^th^ and 75^th^ percentiles from the distribution made of all median PMTHs from the sessions included in the group. Overall are included: P5: N=4, n=8, P6: N=3, n=5; P7: N=5, n=12; P8: N=5, n=8; P9: N=5, n=11; P10: N=1, n=1; P11: N=1, n=1; P12: N=5, n=7 (N: number of mice, n: number of imaging sessions). In all panels, the thin straight gray lines represent the 5^th^ percentile, the median and the 95^th^ percentile of the distribution made of all median PMTHs resulting from surrogate raster plots from the sessions included in the group. Results to build the PMTH as well as CICADA configuration file to reproduce the analysis are available in Figure 2 - Source Data 1. **(B)**. Distribution of post-movement activity across age. Each boxplot is built from all detected movements for the given age group. Whiskers represent the 5^th^ and 95^th^ percentiles with post-movement activity falling above or below represented as small dots. The average post-movement activity observed for each mouse pup is represented by the large dots color coded from light gray (P5) to black (P12). The red area illustrates the movement falling in the category of ‘inhibiting’ movements. Results to build the distributions as well as CICADA configuration file to reproduce the analysis are available in Figure 2 - Source Data 1. **(C)**. Distribution of the proportion of ‘inhibiting’ movements across age. Each dot represents a mouse pup and is color coded from light gray (P5) to black (P12). The open blue circles represent the median of the age group. The red line shows a sigmoïdal fit with V50=9.015, r^2^=0.75. Results to build the distribution as well as CICADA configuration file to reproduce the analysis are available in Figure 2 - Source Data 1. **(D)**. Distribution of the proportion of significatively immobility-associated cells as a function of age. Each dot represents a mouse and is color coded from light gray (P5) to black (P12). The open blue circles represent the median of the age group. Results to build the distribution as well as CICADA configuration file to reproduce the analysis are available in Figure 2 - Source Data 1.

### GABAergic neurons remain activated by spontaneous movement throughout the two first postnatal weeks

As a first step to identify the circuit mechanisms for this switch, we focused on local circuits and disentangled the respective contribution of local GABAergic neurons and principal cells to CA1 dynamics as well as their relation to movement. To this aim, we identified GABAergic neurons with the expression of a red reporter (tdTomato) in *GAD67Cre* pups virally infected with AAV9-FLEX-CAG-tdTomato and AAV1.hSyn.GCaMP6s (**Figure 3A, B**). In addition, we used these imaging experiments to train a cell classifier inferring interneurons in the absence of any reporter with 91 % reliability (Denis et al., 2020). “Labeled” and “inferred” GABAergic neurons were combined into a single group referred to as “interneurons’’ in the following (**Figure 3 - figure supplement 2B, C**). As illustrated in a representative raster plot from a P5 mouse, both pyramidal cells (black) and interneurons (red) were activated during movement (vertical grey lines, **Figure 3A**). This was confirmed when computing the PMTH for pups aged less than P9, with the activation of the two neuronal populations after movement exceeding chance level (P5-8: N=17, n=33, pyramidal cells: baseline value = 0.51 %, peak value = 2.1 %, interneurons: baseline value = 2.1 %, peak value 7.9 %, N: number of mice, n: number of imaging sessions, **Figure 3A**). In line with above results (**Figure 2A**), pups older than P9 showed a significant reduction in the proportion of active pyramidal cells following movement (**Figure 3B**, P10-12: N=7, n=9, baseline value = 1.3 %, trough value = 0.4 %). In contrast, interneurons remained significatively activated following movement even past P9 (**Figure 3B**, P10-12: N=7, n=9, baseline value = 3.9 %, peak value = 10 %). We conclude that the link between movement and activity evolves differentially towards the start of the second postnatal week when comparing pyramidal neurons and GABAergic interneurons, the former being inhibited or detached from movements while the latter remaining activated. This suggests that pyramidal neurons could be directly inhibited by local interneurons after the first postnatal week, following a functional maturation of GABAergic outputs onto principal cells. Alternatively, this could result from differential changes in the synaptic inputs driving both cell types. In the following, we have addressed both, non-mutually exclusive, hypotheses.

**Figure 3:**
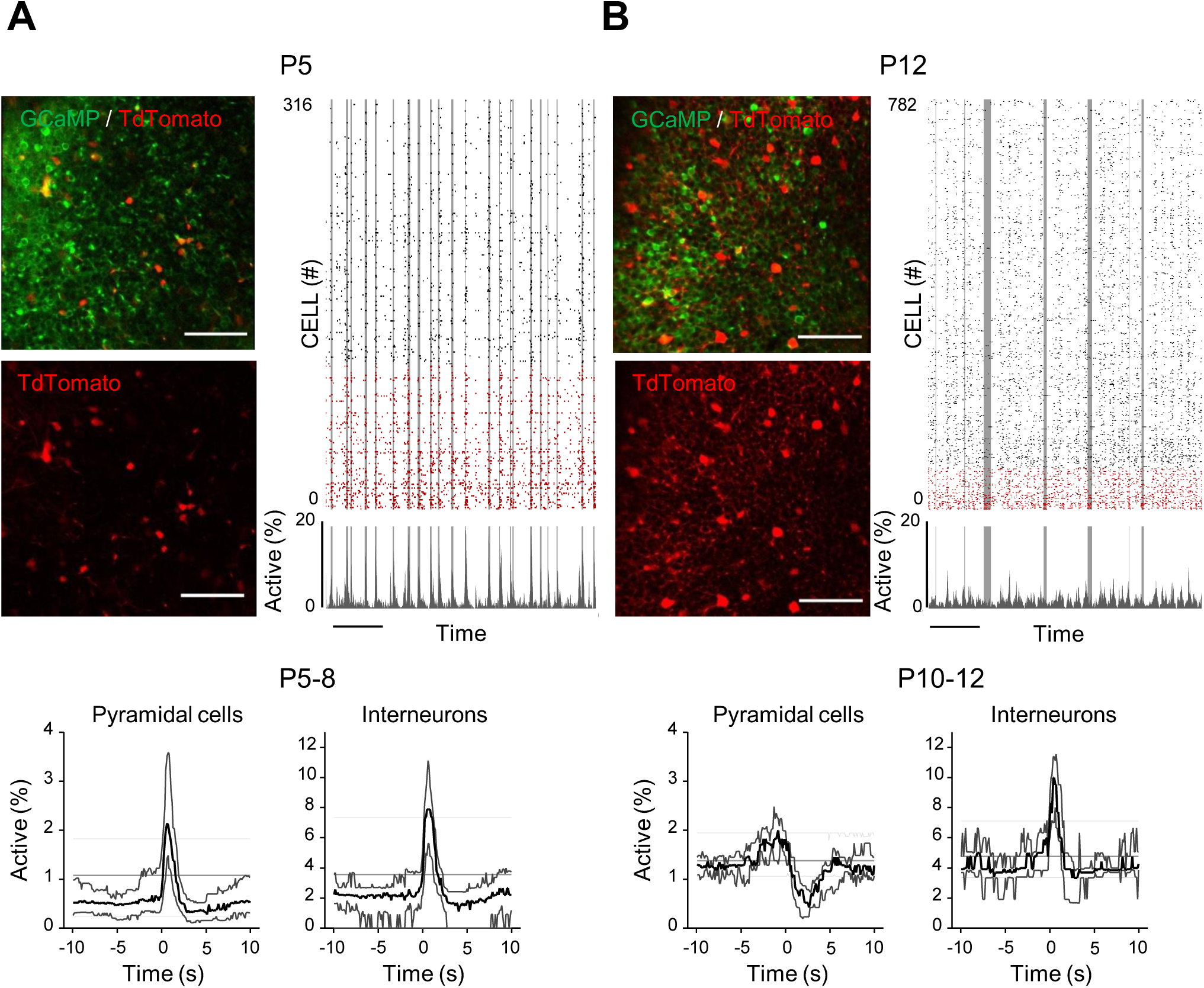
Differential recruitment of CA1 glutamatergic and GABAergic neurons. **(A)**. Top panel: Imaged field of view and associated raster plot from an example imaging session in the *stratum pyramidale* from one P5 *Gad67Cre* mouse pup (scale bar = 100μm). Imaged neurons expressed GCaMP6s. Interneurons were identified by the Cre-dependent expression of the red reporter tdTomato. In the raster plot neurons are sorted according to their identification as pyramidal cells (black) or interneurons (red), vertical gray lines indicate movements of the mouse. Scale bar: 60 seconds. Bottom panel: PMTHs for pyramidal cells and interneurons combining all imaging sessions from mice aged between P5 and P8. The dark line indicates the median value and the thick grey lines represent the 25^th^ and 75^th^ percentiles from the distribution made of all median PMTH obtained from the sessions included in the group. Thin gray lines represent the 5^th^, median and 95^th^ from the distribution made of all median PMTH obtained from surrogate raster plots from the sessions included in the group **(B)**. Same as (A). But illustration is made with one P12 *Gad67Cre* mouse pup and PMTHs are built with all imaging sessions from pups aged between P10 and P12. Note the presence of red labeled processes in the neuropil of the *stratum pyramidale* of P12 in contrast to P5. Results to build the PMTHs as well as CICADA configuration files to reproduce the analysis are available in Figure 3 - Source Data 1.

We first compared the developmental time course of extra-hippocampal synaptic afferences onto CA1 GABAergic neurons and pyramidal cells using a rabies retrograde tracing method (Wickersham et al., 2007). We focused on changes that may occur around the end of the first postnatal week. To do so, two groups were compared, an *early* (AAV1-hSyn-FLEX-nGToG-WPRE3 -Helper virus-injected at P0, SAD-B19-RVdG-mCherry -pseudotyped defective rabies virus- at P5 and immunohistochemistry (IHC) at P9, **Figure 3 - figure supplement 1A-C**) and a *late* one (AAV1-hSyn-FLEX-nGToG-WPRE3 -Helper virus- injected at P0, SAD-B19-RVdG-mCherry -pseudotyped defective rabies virus- at P9, IHC at P13, **Figure 3 - figure supplement 1D-F**). Injections were performed in either *GAD67Cre* or *Emx1Cre* pups in order to specifically target GABAergic or glutamatergic cells, respectively. Four *GAD67Cre* pups (2 early and 2 late injections) and three *Emx1Cre* pups (1 early and 2 late) were analyzed with injection sites restricted to the hippocampus. Starter and retrogradely labeled cells were found all over the ipsilateral hippocampus. For both *GAD67Cre* and *Emx1Cre* pups, we found no striking difference in the retrogradely labeled extra-hippocampal regions between the *early* and *late* groups. In agreement with previous studies (Supèr and Soriano, 1994), we found that GABAergic and glutamatergic neurons in the dorsal hippocampus received mainly external inputs from the entorhinal cortex, the medial septum and the contralateral CA3 area (retrogradely labeled cells in these regions were found in 4 out of 4 *GAD67Cre* pups and 3 out of 3 *Emx1Cre* pups, **Figure 3 - figure supplement 1**). Thus, we could not reveal any major switch in the nature of the extra-hippocampal inputs impinging onto local CA1 neurons. Thus, we next explored the maturation of local somatic GABAergic innervation given its significant evolution throughout that period (Jiang et al., 2001; Marty et al., 2002; Morozov and Freund, 2003) as well as our observation of a dense tdTomato signal in the pyramidal layer from *GAD67Cre* mouse pups at P12 (**Figure 3B**), not visible at P5 (**Figure 3A**).

### Abrupt emergence of a functional somatic GABAergic innervation at the beginning of the second postnatal week

We first analyzed the anatomical development of somatic GABAergic innervation within the CA1 pyramidal layer from P3 to P11, focusing on the innervation from putative parvalbumin-expressing Basket Cells (PVBCs), its main contributor. To this aim, we performed immunohistochemistry against Synaptotagmin2 (Syt2) which has been described as a reliable marker for parvalbumin positive inhibitory boutons in cortical areas (**Figure 4A**, (Sommeijer and Levelt, 2012)). Using a custom-made Fiji-plugin (RINGO, see methods), we quantified the surface of the pyramidal cell layer covered by Syt2 labeling at different stages and found that between P3 and P7, PV innervation remained stable (median values: P3: 0.34 %, P5: 0.57 %, P7: 0.49 %, 3 mice per group, **Figure 4B**). However, around P9, a sudden increase in the density of positive labelling was observed (P9: 1.03 %, P11: 1.48 %, 3 mice per group, **Figure 4B**). These results are consistent with previous work (Jiang et al., 2001; Marty et al., 2002), as well as with our tdTomato labeling (**Figure 3A-B**). They also match the transition observed in CA1 dynamics (**Figure 3**). We next tested whether GABAergic axons in the pyramidal layer were active during periods of movement. In order to increase success rate for imaging axons in the pyramidal layer, we restricted these experiments to P9-10, i.e., when axons start densely innervating the CA1 layer. To do so, we restrained the expression of the calcium indicator GCaMP6s to the axon (Broussard et al., 2018) of interneurons using *GAD67Cre* mouse pups and specifically imaged axonal arborisation in the pyramidal cell layer (**Figure 4C, left panel**). Fluorescence signals were extracted from axonal branches using PyAmnesia (a method to segment axons, see methods, **Figure 4C, right panel)**, and then normalized using z-score (see methods). As expected, (see **Figure 3B)**, an increase in the fluorescent signal from GABAergic axonal branches was observed following movement (P9-10: n=3 mice, **Figure 4D**). As a result, we reasoned that an increase in perisomatic GABAergic inhibition could contribute to the reduction of activity observed after movement during the second postnatal week in pyramidal neurons.

**Figure 4:**
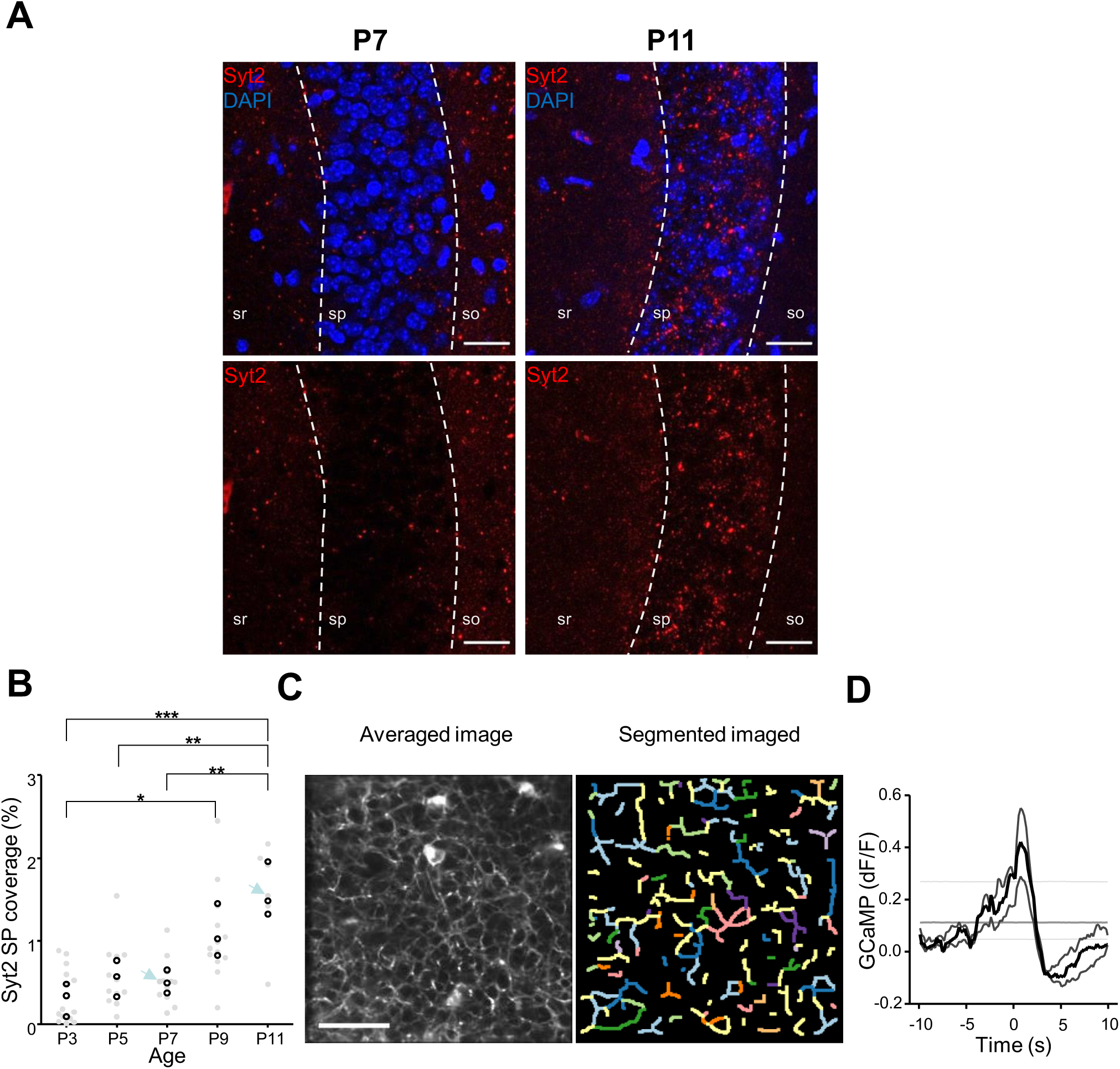
Emergence of perisomatic GABAergic innervation. **(A)**. Representative example confocal images of the CA1 region in a P7 (left) and a P11 (right) mouse pup. DAPI staining was used to delineate the *stratum pyramidale* (sp) from the *stratum radiatum* (sr) and *stratum oriens* (so, top row). Synaptotagmin-2 labeling (Syt2) is shown in the top and bottom rows. Illustrated examples are indicated by red dots in the associated quantification in (B). Scale bar= 20 μm. **(B)**. Fraction of the pyramidal cell layer covered by Syt2 positive labeling as a function of age. Each gray dot represents the average percentage of coverage from two images taken in the CA1 region of a hippocampal slice. Open black dots are the average values across brain slices from one mouse pup. Blue arrows indicate the slices used for illustration in (A). A significant effect of age was detected (One-Way Anova, F=13.11, p=0.0005). Multiple comparison test shows a significant difference between age groups (Bonferroni’s test, *: p<0.05, **: p<0.01, ***: p <0.001). **(C)**. Averaged image of a field of view in the pyramidal cell layer of a P9 *GAD67Cre* mouse pup injected with a Cre-dependent Axon-GCaMP6s indicator (left) and the segmented image resulting from PyAmnesia (right). **(D)**. PMTH showing the df/f signal centered on the onsets of animal movement (N=3, n=3). The dark gray line indicates the median value, and the light gray lines represent 25^th^ and 75^th^ percentile. Results obtained from surrogates are represented by light gray lines.

### Increasing feedback inhibition in two-population models reproduces the developmental transition

To test whether an increase in perisomatic inhibition alone can explain the switch in network dynamics between the first and second postnatal weeks, we simulated a two-population network mimicking the development of perisomatic innervation (**Figure 5A**, see methods). Using a rate model and a Leaky Integrate and Fire (LIF) model, we show that increasing the strength of perisomatic inhibition can account for the experimentally observed decrease of responses to movement-like feedforward inputs (**Figure 5B**). Durations of feed-forward inputs were chosen similar to experimental movement durations (see **Figure 5 - figure supplementary 1A** for a log-normal of the movement durations). When inhibition is weak, the average activity of the pyramidal neurons increases at the onset of a given twitch. Then, it quickly relaxes to the baseline with a timescale that follows the synaptic time constant (**Figure 5B**, left panel). In the presence of strong inhibition, there is a reduction in response to movement inputs. In addition, due to strong feedback inhibition following the movement responses, network activity relaxes to the baseline with an undershoot (**Figure 5B**, right panel), recapitulating the experimental findings (see **Figure 2A**). Similar PMTHs were obtained for interneurons (see **Figure 5 - figure supplementary 1 B**).

**Figure 5:**
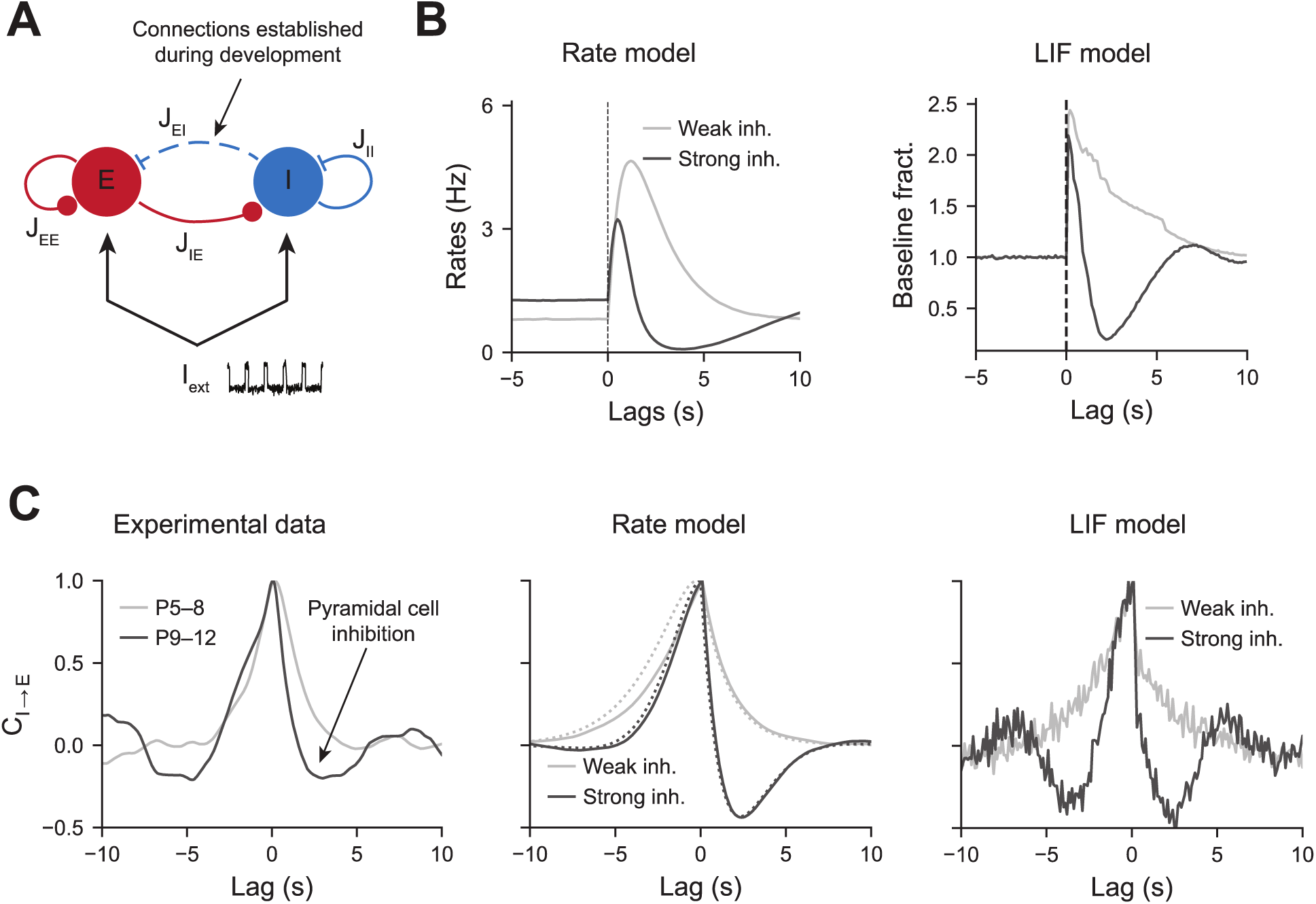
Modeling the effects of perisomatic inhibition on pyramidal cell response. **(A)**. The model consists of two populations, Excitatory (E) and Inhibitory (I) receiving feedforward input I_ext_. The interaction strengths, Jab represent the effect of the activity of population b on a. We study the effects of perisomatic inhibition on the activity of pyramidal cells by varying the parameter J_EI_. (**B)**. PMTH, response of excitatory neurons to pulse input in the rate model and LIF network. **(C)**. Cross-correlations during periods of immobility, in experimental data (left), rate model (middle), and LIF network (right). In the rate model, dotted lines are the predicted correlation from the analytic expressions and solid lines are the results from numerical integration.

To further show the importance of perisomatic inhibition on network dynamics, we computed the cross-correlogram between interneurons activity and pyramidal cell activity in the absence of movement (P5-8 light gray curve, P9-12 dark curve, **Figure 5C, left panel**). For P9-12, we observe a rapid drop in the correlation at positive time which means that elevated inhibitory activity is followed by a strong decrease in excitatory activity. Notice that this drop is absent for P5-8, where the cross-correlogram is mostly symmetric around t=0. Both the rate and LIF models displayed similar activity correlograms as the experimental data, with an undershoot in the presence of strong inhibitory feedback for positive time. We showed through numerical simulations that similar changes could be observed in a more realistic spiking network model (**Figure 5C, right panel**). Auto-correlograms of the excitatory and inhibitory activity were also measured and compared to our model predictions (see **Figure 5 - figure supplementary 1C**).

The consistency of our models with the experimental cross-correlograms, which were computed from the activity recorded during periods of immobility, further shows that the observed network dynamics and, in particular, the correlation undershoots most likely result from recurrent perisomatic inhibition rather than a feedforward drive from upstream areas. Therefore, in our model, the maturation of perisomatic inhibition alone was sufficient to support a switch in network dynamics.

## DISCUSSION

Using for the first time *in vivo* two-photon calcium imaging in the hippocampus of non-anesthetized mouse pups and a deep-learning based approach to infer the activity of principal cells and interneurons, we show that the end of the first postnatal week marks a salient step in the anatomical and functional development of the CA1 region. Indeed, within two days (P8-10), the link between CA1 principal cells activity and self-triggered movements is inverted and neurons are preferentially active during immobility periods. This is likely due to the time-locked anatomical and functional rise of somatic GABAergic activity given that interneurons remain highly active throughout this period, including in response to spontaneous movements. In this way, CA1 circuits start detaching from external inputs. Given the importance of internal dynamics for hippocampal function and cortical circuits operation in general, this is likely to be a critical general step in the proper maturation of cognitive circuits.

### Early postnatal calcium activity in CA1 is driven by sensorimotor inputs

We found that, until P7-9, spontaneous movements are followed by a significant peak in calcium events in the CA1 principal cell layer and that most neuronal activity occurs during synchronous calcium events. This early link between sensorimotor inputs and early cortical dynamics has been previously reported using electrophysiological recordings in various areas and species, including humans (Milh et al., 2006). Here we extend that observation to calcium transients, which not only indirectly report action-potential firing as well as other modes of cell activation during development but also critically regulate activity-dependent genetic processes. In addition, we could describe the response to these movements with single-cell resolution. It is possible that other movements, like whisker movements, that we have not detected also contribute to the patterning of CA1 activity. It is also possible that self-generated activity from other sensory organs but independent from movement, like the retina or the olfactory bulb, also contribute to hippocampal dynamics, the same way they are relayed to sensory cortices. Interestingly, in contrast to previous reports (Tiriac et al., 2014), we did not observe any significant difference between twitches (occurring mainly during active sleep) and longer, more complex movements. This may reveal a difference between calcium imaging and electrophysiology, the former sampling from a larger population but at a lower temporal and spike signal resolution. The patterning of CA1 dynamics in the large imaged population did not reveal any obvious spatial distribution for movement-activated cells but we cannot exclude that these would vary along the radial and transverse directions, which are the two main axes of principal cell development (Caviness, 1973), and are differentially targeted by perisomatic PV basket cells (Lee et al., 2014; Valero et al., 2015).

Passed the end of the first postnatal week, between P8 and P10, a significant decrease in the fraction of coactive principal cells following movement was observed (while interneurons remained mostly activated by movement). We cannot exclude that some spikes fell below the threshold for detection of calcium events. In this case, rather than full inhibition, it may be that a strong shortening of the time window for neuronal integration occurred (due to feedback inhibition), which would limit the number of spikes produced by principal cells and thus keep them below detection levels. Yet, a novel machine-learning based algorithm (Denis et al., 2020) was used since it was especially designed to infer activity in the dense CA1 pyramidal cell layer. This change in the polarity of principal cells’ response to movements is quite abrupt as it happens within less than two days (between P8 and 10). This contrasts with the progressive evolution of single cell firing frequencies but matches the fast redistribution of neuronal firing towards immobility periods. In this way, hippocampal neuronal dynamics “internalize” as they stop being driven by movements and preferentially occur within rest.

This “internalization” of hippocampal dynamics is reminiscent of similar phenomena observed in other cortical areas, such as the barrel cortex where whisker stimulation induces a reduction in the size of cell assemblies following P9 while the same stimulation widens cell assembly size a few days before (Mòdol et al., 2019). It is also reminiscent of the recently described transient quiescent period observed in the somatosensory cortex using extracellular electrophysiological recordings (Dominguez et al., 2021). Last, it goes in hand with a sparsification of activity, which is a general developmental process supported by the emergence of inhibition (Golshani et al., 2009; Rochefort et al., 2009; Wolfe et al., 2010).

### Circuit basis for the movement-triggered inhibition of CA1 dynamics

Our results demonstrate that the “internalization” of CA1 dynamics occurring at the end of the first postnatal week most likely relies on structural changes in local CA1 circuits rather than rewiring of the long-range extrahippocampal connectivity.

The long-range circuits mediating the bottom-up flow of self-triggered or externally generated sensory information to the hippocampus are starting to be elucidated. The two main structures directly transmitting sensorimotor information to the dorsal CA1 are the entorhinal cortex and septum. The former processes multisensory information from all sensory cortices (visual, auditory, olfactory, somatosensory), including movement-related sensory feedback (Rio-Bermudez and Blumberg, 2021)and was shown to be activated by spontaneous twitches prior to CA1 (Mohns and Blumberg, 2010; Rio-Bermudez et al., 2020; Valeeva et al., 2019a) while the latter is more likely to be involved in transmitting internal information (Fuhrmann et al., 2015; Wang et al., 2015) as well as unexpected environmental stimuli (Zhang et al., 2018). In addition to these two canonical pathways, one cannot exclude the involvement of a direct connection from the brainstem, given their existence in the adult and their role in promoting sleep as well as motor twitches (Liu et al., 2017; Szőnyi et al., 2019). However, our retrograde tracing experiments did not reveal any direct connection between the CA1 cells and the brainstem at the early ages analyzed here. In addition, we found that both CA1 interneurons and principal cells receive inputs from the septum and entorhinal cortex before the time of the switch (i.e. P9) and that there was no major qualitative change of inputs after, as expected from previous work (Supèr and Soriano, 1994). Still, these experiments do not allow a quantitative assessment of the number of inputs nor the type of inputs (GABAergic, cholinergic, etc.) and we cannot fully exclude that a stronger, or different source of excitatory drive would be impinging onto interneurons after the switch. Therefore, future optogenetic and slice physiology work is needed to characterize the bottom-up information flow onto specific components of the local CA1 circuits. Similarly, one cannot exclude a change in the CA3 to CA1 connectivity. Indeed, Schaffer collaterals are known to reach CA1 roughly around the end of the first postnatal week (Durand et al., 1996). In addition, roughly at the time of the switch, do we see the emergence of SWRs (Buhl and Buzsaki, 2005), a pattern strongly relying on CA3 inputs and perisomatic GABAergic transmission. However, we could not restrict the pool of starter cells to the CA1 region in our retrograde viral tracing experiments, which precluded analysis of the development of CA3-CA1 connectivity. Interestingly, among the external inputs onto CA1 described above, the entorhinal cortex and CA3 were both shown to exert a mild influence on the organization of intrinsic CA1 dynamics, possibly pointing at a critical role of local interneurons in this process (Zutshi et al., 2021).

As indicated by our computational model, the disengagement from movement of CA1 dynamics can be fully explained by the observed rise in anatomical (Syt2 labeling) and functional (axonal GCaMP imaging) connectivity from perisomatic GABAergic cells onto pyramidal cells at the onset of the second postnatal week. This increased connectivity could not be easily captured with our retrograde viral labeling since the absence of early PV expression precludes the identification of PV basket cells, the most prominent subtype of perisomatic GABAergic cells, among retrogradely labelled cells in *Emx1Cre* pups. Early anatomical studies had already indicated that an increase of somatic GABAergic inhibition, including from CCK-basket cells, occurred in CA1 during the first postnatal week (Danglot et al., 2006; Jiang et al., 2001; Marty et al., 2002; Morozov and Freund, 2003). However, this rise was expected to be more progressive and not as abrupt as observed here, as it happened within two days. If the axonal coverage of the stratum pyramidale by PV-basket cells axons increases, we cannot exclude that this is a general phenomenon, concerning all perisomatic subtypes, including soma-targeting CCK-expressing basket cells which develop anatomically at around the same time (Morozov and Freund, 2003) or chandelier cells. In addition, our computational model indicates that the emergence of feed-back inhibition is sufficient to reproduce the developmental shift observed here, which could also involve other types of CA1 interneurons, including dendrite-targeting ones.

Interestingly, a similar rise of somatic GABAergic axonal coverage occurs in the barrel cortex at the same time. Indeed, recent connectomic mapping using 3D-electron microscopy in that region revealed that the preferential targeting of cell bodies by GABAergic synapses increased almost threefold between postnatal days 7 and 9 (Gour et al., 2020), whereas two-photon imaging of putative GABAergic somatic axons in the same region revealed broader domains of co-activation (Mòdol et al., 2019). This time period for the shift may be synchronous within brain regions involved in sensorimotor integration such as the hippocampus and somatosensory cortex. Otherwise, PV expression was shown to develop sequentially in a region-specific manner (Reh et al., 2020) following their intrinsic developmental age (Donato et al., 2017).

We found that many principal cells are inhibited by movement while most imaged GABAergic cells remained activated during the second postnatal week. This therefore indirectly suggests a net inhibitory effect of GABAergic transmission after the first postnatal week. This is expected since the shift from excitatory to inhibitory synaptic transmission was reported to occur earlier in the hippocampus *in vivo* (Murata and Colonnese, 2020). On a side note, the lack of somatic GABAergic inputs before P7 indicates that the early excitatory GABAergic drive in CA1 circuits likely originates from non-somatic GABAergic interneurons, which include long-range, dendrite-targeting or interneuron-specific interneurons. The circuit role of excitatory GABAergic transmission should be revisited taking into account this new finding.

The movement-associated inhibition can result equally from feedforward (direct activation from movement-transmitting inputs such as the entorhinal cortex) or feedback (from local CA1 cells) inhibition. Our experiments do not allow disentangling both circuits. A progressive strengthening of feedback inhibition circuits through the strengthening of local CA1 principal cells inputs onto perisomatic GABAergic neurons is also possible, as it occurs in the developing somatosensory cortex (Anastasiades and Butt, 2012). The inhibition of activity following movement is likely to be occurring during a transient developmental period. Indeed, in the adult, both interneurons and principal cells usually increase their activity as the animal moves (Fuhrmann et al., 2015). Therefore the switch observed here opens another developmental time-window that probably closes with the emergence of perineuronal nets and cell activation sequences at the end of the third developmental week (Farooq and Dragoi, 2019; Horii-Hayashi et al., 2015; Muessig et al., 2019). We would like to propose this developmental window, to be the critical period for CA1 development, a period during which experience-dependent plasticity can be observed.

## Conclusion

Cognitive hippocampal maps rely on two forms of representation, one that is map-based or allocentric and the other that is self-referenced, or egocentric and requires body movement. We would like to propose that the early postnatal period described here, where the hippocampus learns the statistics of the body, and which terminates with the rise of a recurrent inhibitory network is a key step for the emergence of an internal model onto which exploration of the external world can be grafted. An imbalance between internal and environmental hippocampal representations due to a miswiring of local somatic inhibition could have major outcomes. It could be at the basis of several neurodevelopmental disorders, including autism spectrum disorders (ASD) and schizophrenia. Interestingly, both disorders have been associated with an aberrant maturation of PV-expressing interneurons (Gogolla et al., 2014; Jurgensen and Castillo, 2015; Lewis et al., 2005). In addition, the proper development of the peripheral sensory system, which is partly initiating the early CA1 dynamics reported in our study, is also critically involved in ASD (Orefice et al., 2016). The period described here corresponds to the third trimester of gestation and likely extends postnatally given the protracted integration of GABAergic interneurons into functional circuits in the human brain (Murphy et al., 2005; Paredes et al., 2016). Future work should determine when a similar rise in somatic inhibition occurs in human infants and test whether it could constitute a valuable biomarker for cognitive neurodevelopmental disorders.

---

**Figure 1 - figure supplement 1.**
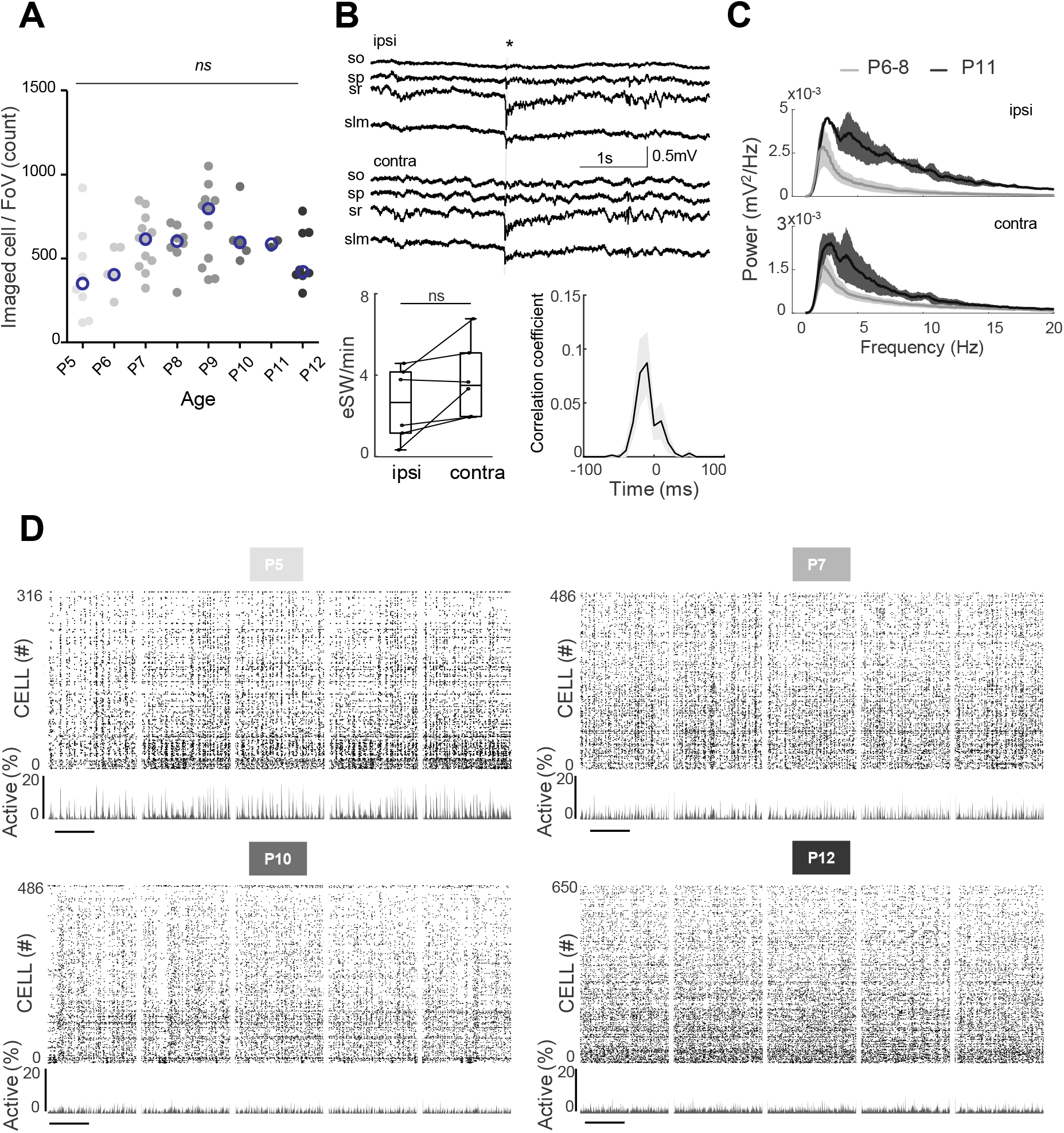
**(A)**. Distribution of the number of cells imaged in each field of view as a function of age. Each dot represents a field view and is color coded from light gray (P5) to black (P12). The open blue circles represent the median of each group. No effect of age was observed on the number of imaged cells. **(B)**. Top panels: Example of the electrophysiological recording done in the ipsi (hippocampus with window implant) and contralateral (intact hippocampus) hippocampi. Use of multisite silicon probe allowed simultaneous recordings in different hippocampal layers (strati oriens (so), pyramidale (sp), radiatum (sr) and lacunosum moleculare (slm)). The eSW is marked by an asterisk. Bottom panels: box plot displays the eSW occurrence. Each dot represents the eSW rate per minute in a mouse pup (N=6). Box plots show median, the bottom and top edges of the box indicate the 25th and 75th percentiles, the whiskers extend to the most extreme data points not considered outliers. Bottom right panel: Cooccurrence of eSWs detected in both hippocampi. **(C)**. Developmental changes in power spectral density of the recorded LFP in the ipsi (hippocampus with window implant) and contralateral (intact hippocampus) hippocampi. Gray and black lines are the averages between the animals of P6-8 and P11 age ranges, respectively. Shading shows jackknife standard deviation. Note, presence of peak in theta frequency band (4-7 Hz) in P11 animals. **(D)**. Raster plots from the imaging sessions used for illustration in Figure 1A, showing all cells over the full recording, scale bar is 2 minutes.

**Figure 2 - figure supplement 1.**
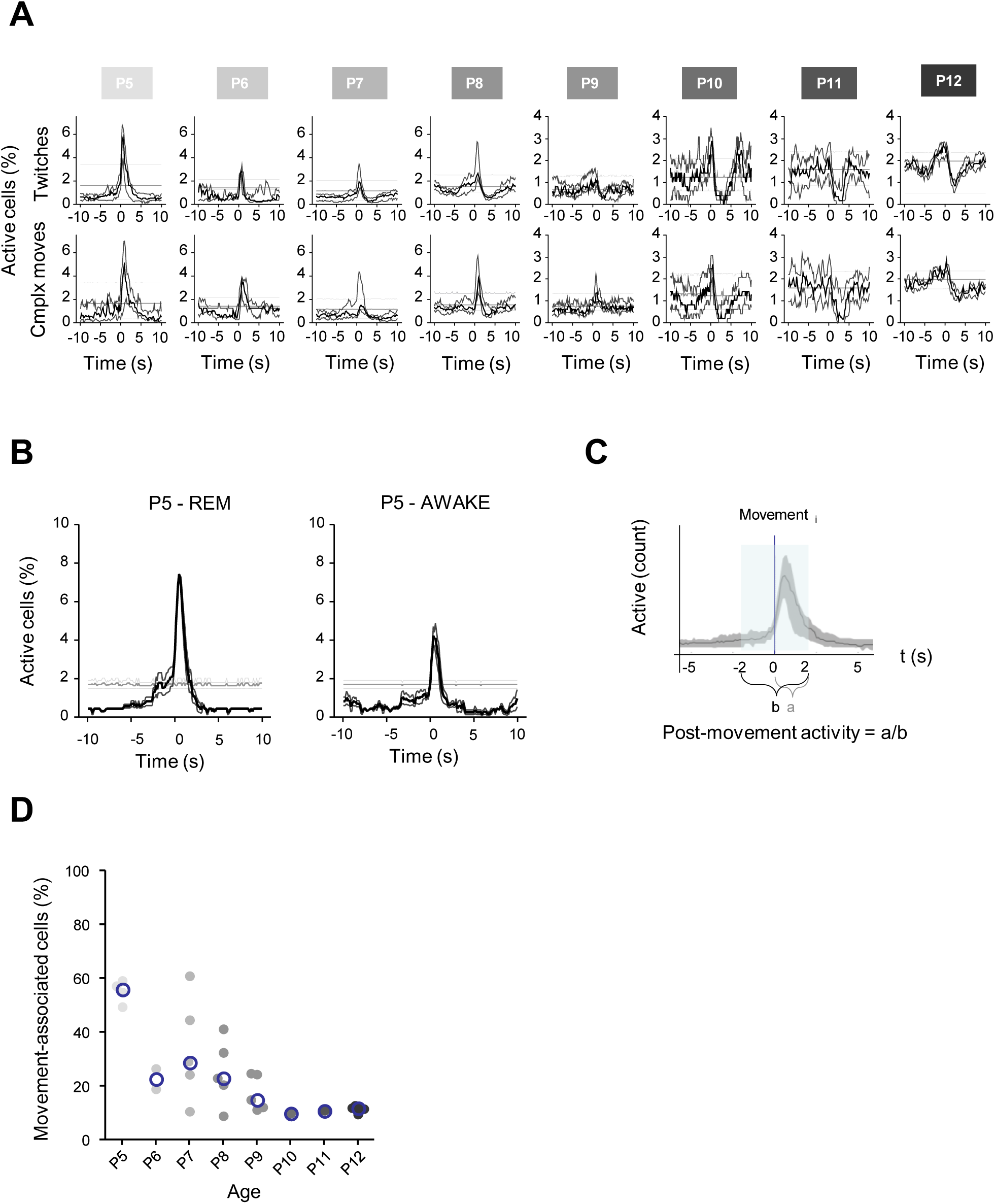
**(A)**. PMTHs for each age group built on twitches only (top row) or on complexe movements only (bottom row). Procedure to group sessions is similar to the one described in Figure 2 **(B)**. PMTHs from one P5 mouse pup (including 2 imaging sessions) combining all detected movements during REM sleep (‘P5 -REM’) or all detected movement during Wakefulness (‘P5 - AWAKE’). **(C)**. Definition of the post-movement activity. **(D)**. Distribution of the proportion of cells significatively associated with movement as a function of age. Each dot represents a mouse pup and is color coded from light gray (P5) to black (P12). The open blue circles represent the median for each group.

**Figure 3 - figure supplement 1.**
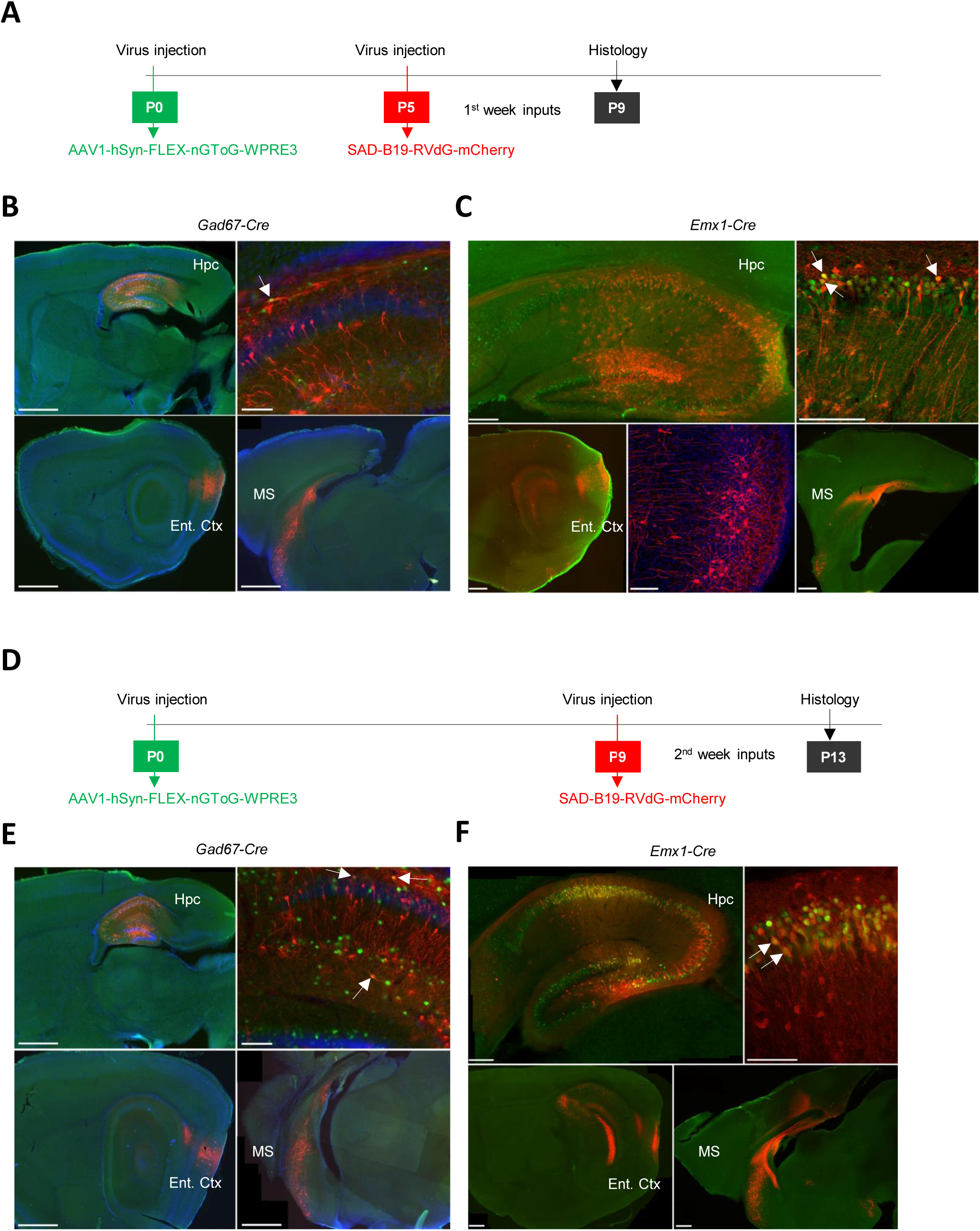
**(A)**. Experimental timeline used to investigate the inputs received by pyramidal cells (*Emx1Cre* mice) and interneurons (*GAD67Cre* mice) during the first postnatal week (early group). At P0 and P5 we injected the helper virus AAV1-hSyn-FLEX-nGToG-WPRE3 and the rabies SAD-B19-RVdG-mCherry into the hippocampus of *GAD67Cre* (Figure 3B) mice and *Emx1Cre* mice (Figure 3C) respectively. Brains are next collected for histological/anatomical analysis at P9. **(B)**. Confocal images from early group *GAD67Cre* mice showing the injection sites in the dorsal hippocampus on sagittal slices at lower magnification (top left, scale bar= 1 mm), and higher magnification (top right, scale bar=100 μm). Primo infected cells are labelled in green, the starter cell indicated by the arrow shows co-expression of the helper virus (green) and rabies virus (red). Presynaptic cells, labelled only in red, are localized in the entorhinal cortex (bottom left, scale bar= 1 mm) and in the medial septum (bottom right, scale bar= 1 mm). Hpc= Hippocampus; Ent. Ctx= Entorhinal Cortex; MS= Medial Septum **(C)**. same as in (B) but for *Emx1Cre* mice. Scale bars from top left to bottom right: 200 μm, 100 μm, 500μm, 100 μm and 500 μm. Middle bottom image is a higher magnification of the bottom left panel. **(D)**. Same as in (A) but for the late group (P9-P13). **(E)**. Same as in (B) but for the late group. Scale bars from top left to bottom right: 1 mm, 100 μm, 1 mm and 1mm. **(F)**. Same as in (C) but for the late group. Scale bars from top left to bottom right: 200 μm, 100 μm, 500μm, and 500 μm.

**Figure 3 - figure supplement 2.**
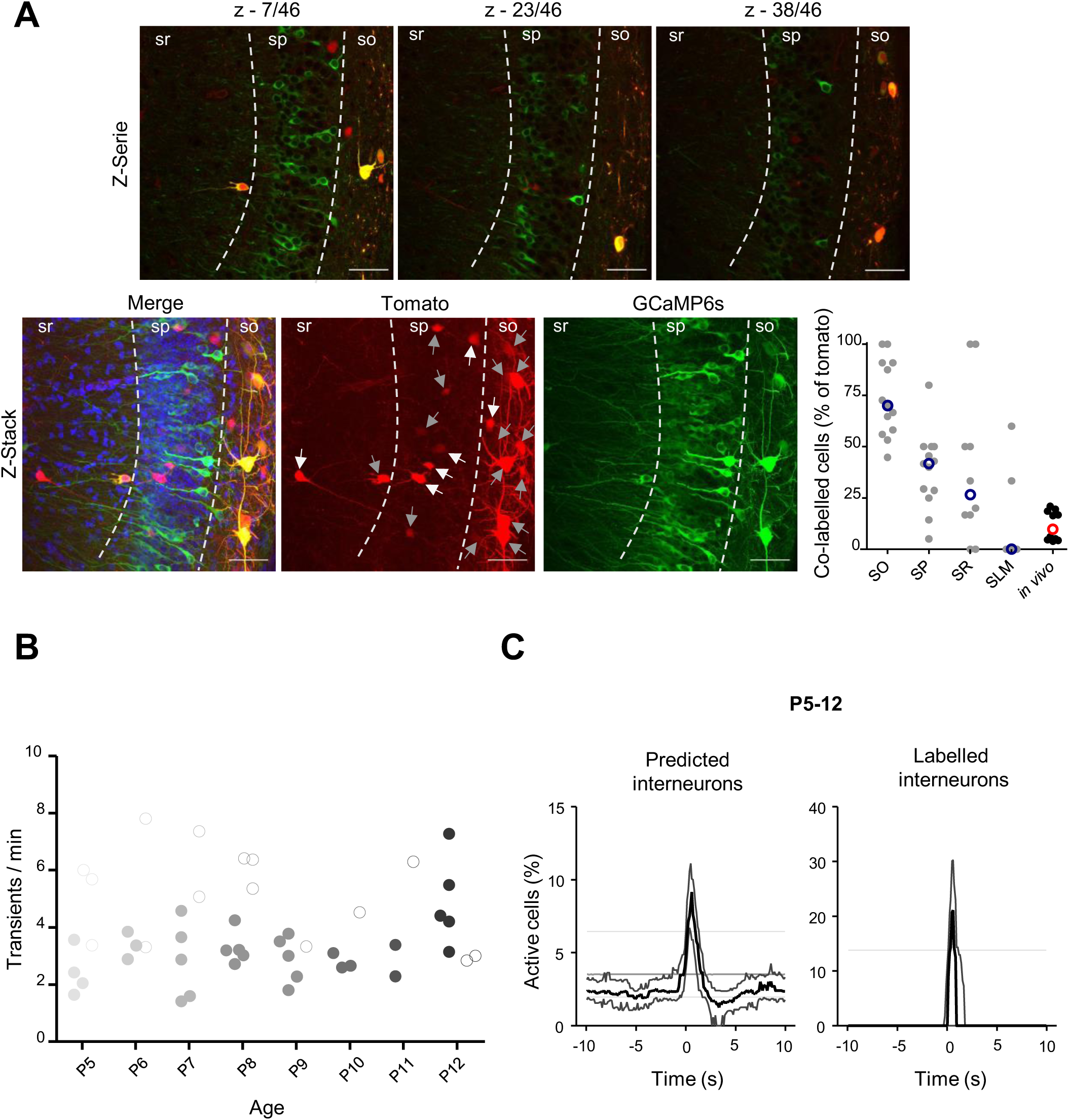
**(A)**. Top row: confocal images taken from a 69 μm z-stack (z-step: 1.5 μm) of a brain slice from a *Gad67Cre* mouse pup injected with both AAV9-FLEX-tdTomato and AAV1.hSyn.GCaMP6s showing the co-infection between tdTomato and GCaMP6s in hippocampal interneurons. Bottom row: maximum intensity projections of the example z-stack showed in (A). On the maximal intensity projection from tdTomato signal white arrows indicate interneurons not expressing GCaMP6s, gray arrows indicate interneurons expressing GCaMP6s. The proportion of tdTomato cells expressing GCaMP6s is shown on the plot for each layer of the region CA1 of the hippocampus. The proportion of tdTomato cells expressing GCaMP6s *in vivo* not only take into account the co-expression of the two but represent the proportion of tdTomato cells having a GCaMP6s signal that was not classified as ‘noise’ explaining the relatively low proportion in comparison with the observed co-infection rate. Scale bars= 50 μm **(B)**. Evolution of the number of transient per minute observed in ‘labelled’ interneurons and ‘inferred’ interneurons from P5 to P12. Each dot represents the average obtained from one mouse and is color coded from light gray (P5) to black (P12). Filled dots represent ‘inferred’ interneurons, open dots represent ‘labelled’ interneurons. **(C)**. PMTHs combined all imaging sessions between P5 and P12 showing the activation after movement of both ‘labelled’ and ‘inferred’ interneurons.

**Figure 5 – figure supplement 1.**
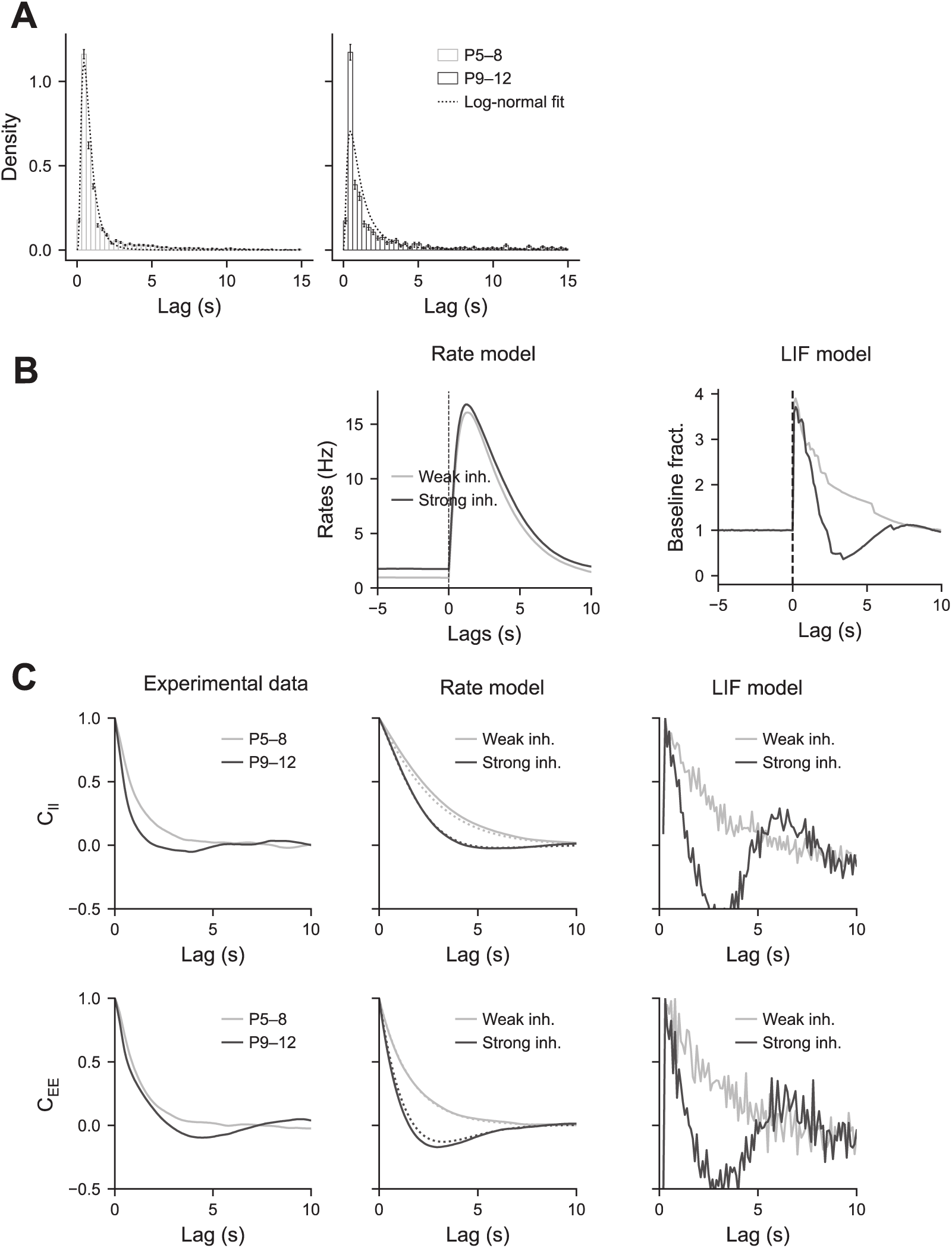
**(A)**. Movement duration histograms. These histograms were fit by log-normal distributions. **(B)**. PMTHs of the interneurons predicted by the rate and LIF models. **(C)**. Auto-correlograms of the activity for the pyramidal cells and interneurons.

**Figure 5 – Supplementary table 1:** Details of the parameters used in the rate and LIF models

| Parameter | Symbol | Value |
| --- | --- | --- |
| **Rate model** |  |  |
| Excitatory time constant | $\tau_E$ | 0.7s |
| Inhibitory time constant | $\tau_I$ | 3.0s |
| Connectivity matrix |  |  |
| | $J_{EE}$ | 0.8 |
| | $J_{IE}$ | 0.5 |
| | $J_{II}$ | -0.2 |
| | $J_{EI}^{weak}$ | -0.0 |
| | $J_{EI}^{strong}$ | -0.4 |
| Noise amplitude | $\eta$ | $0.1s^{-1/2}$ |
| External input |  |  |
| | $H_E$ | 0.5 |
| | $H_I$ | 0.1 |
| External drive increase | $\alpha$ | 3.0 |
| Twitch amplitude |  |  |
| | $J_E$ | 7.0 |
| | $J_I$ | 25.0 |
| **LIF model** |  |  |
| Exc. synapse time constant | $\tau_E$ | 1.0s |
| Inh. synapse time constant | $\tau_I$ | 3.0s |
| Membrane time constant | $\tau_m$ | 20.0 ms |
| Threshold potential | $V_{thr}$ | 20.0 mV |
| Reset potential | $V_{reset}$ | 0.0mV |
| Connectivity |  |  |
| Nb. of excitatory neurons | $N_E$ | 512 |
| Nb. of inhibitory neurons | $N_I$ | 128 |
| Connections from exc. neurons | $K_E$ | 32 |
| Connections from inh. neurons | $K_I$ | 16 |
| Connectivity matrix |  |  |
| | $J_{EE}$ | 0.5 |
| | $J_{IE}$ | 2.0 |
| | $J_{II}$ | -0.3 |
| | $J_{EI}^{weak}$ | -0.05 |
| | $J_{EI}^{strong}$ | -0.8 |
| Noise amplitude | $\eta$ | $1.0s^{-1/2}$ |
| External input |  |  |
| | $H_E$ | 1.0 |
| | $H_I$ | 0.1 |
| Twitch amplitude |  |  |
| | $J_E$ | 2.0 |
| | $J_I$ | 10.0 |
| Twitch duration (lognormal) |  |  |
| Mean parameter | $\mu$ | -0.2 |
| Std parameter | $\sigma$ | 0.8 |

## SUPPLEMENTARY MOVIES

***Supplementary Movies 1-3:*** Three examples of calcium imaging movies from P5 mouse pups centered on the onset of a twitch. The twitch is indicated by T on the upper left corner of the movie. Imaging 2 times speed up.

***Supplementary Movies 4-6:*** Three examples of calcium imaging movies from P12 mouse pups centered on the onset of a complex movement. The complex movement is indicated by M on the upper left corner of the movie. Imaging 2 times speed up.

## SOURCE DATA

***Figure 1 - Source Data 1***. Contains 3 folders. **1A**: contains the 4 raw ‘.tiff’ files illustrating the field of views and the CICADA configuration files necessary to plot the contours map and raster plots used for the illustration. **1B**: contain (in an ‘.xlsx’ file) the numerical data used to plot the evolution of the transient in SCE and the CICADA configuration file necessary to reproduce the analysis. **1C**: contain (in an ‘.xlsx’ file) the numerical data used to plot the evolution of the transient per minute and the CICADA configuration file necessary to reproduce the analysis.

***Figure 2 - Source Data 1***. Contains 4 folders. **2A**: contain (in an ‘.xlsx’ file) the numerical data used to plot all the PMTHs (in Figure 2A) and the CICADA configuration file necessary to reproduce the analysis. **2B**: contain (in an ‘.xlsx’ file) the numerical data used to plot Figure 2B and the CICADA configuration file necessary to reproduce the analysis. **2C**: contain (in an ‘.xlsx’ file) the numerical data used to plot Figure 2C and the CICADA configuration file necessary to reproduce the analysis. **2D**: contain (in an ‘.xlsx’ file) the numerical data used to plot Figure 2D and the CICADA configuration file necessary to reproduce the analysis.

***Figure 3 - Source Data 1***. Contains 3 folders. **‘example_FoVs’:** contains the ‘.tiff’ images used for illustration in Figure 3. **‘example_raster_plots’**: contains the 2 CICADA configuration files necessary to reproduce the raster plots used for illustration in Figure 3. **‘psths’**: contains (in an ‘.xlsx’ file) the numerical data used to plot Figure 3 PMTHs and the CICADA configuration file necessary to reproduce the analysis.

***Figure 4 - Source Data 1***. Contains 4 folders. **4A**: contains ‘.tiff’ images used for illustration in Figure 4A. **4B**: contains the numerical data of the plot in Figure 4B. **4C**: contains the CICADA configuration file necessary to plot the contour map shown in Figure 4C. **4D**: contains (in an ‘.xlsx’ file) the numerical data used to plot Figure 4C PMTH and the CICADA configuration file necessary to reproduce the analysis.

## MATERIALS AND METHODS

### MICE

All experiments were performed under the guidelines of the French National Ethics Committee for Sciences and Health report on “Ethical Principles for Animal Experimentation” in agreement with the European Community Directive 86/609/EEC (Apafis#18-185 and #30-959).

### EXPERIMENTAL PROCEDURES AND DATA ACQUISITION

#### Viruses

*In vivo* calcium imaging experiments were performed using AAV1-hSyn-GCaMP6s.WPRE.SV40 (pAAV.Syn.GCaMP6s.WPRE.SV40 was a gift from Douglas Kim & GENIE Project (Addgene viral prep #100843-AAV1; http://n2t.net/addgene:100843; RRID:Addgene_100843)), AAV9-FLEX-CAG-tdTomato (pAAV-FLEX-tdTomato was a gift from Edward Boyden (Addgene viral prep # 28306-AAV9 ; http://n2t.net/addgene:28306 ; RRID:Addgene_28306)), AAV9-hSyn-FLEX-axon-GCaMP6s (pAAV-hSynapsin1-FLEx-axon-GCaMP6s was a gift from Lin Tian (Addgene viral prep # 112010-AAV9 ; http://n2t.net/addgene:112010 ; RRID:Addgene_112010)). Retrograde tracing experiments were performed using AAV1-hSyn-FLEX-nGToG-WPRE3 (Charité #BA-096) and SAD-B19-RVdG-mCherry (Gift from the Conzelmann laboratory).

#### Intracerebroventricular injection

This injection protocol was adapted from already published methods (Kim et al., 2014). Mouse pups were anesthetized on ice for 3 – 4 min, and 2 μL of viral solution (titration at least 1 × 10^13^ vg / mL) were injected in the left lateral ventricle whose coordinates were estimated at the 2/5 of the imaginary line between the lambda and the eye at a depth of 0.4 mm. Correct injection was visualized by the spreading of the viral-dye mixture (1/20 of fast blue). In SWISS mouse pups we injected 2 μL of AAV2.1-hSyn-GCAMP6s.WPRE.SV40, in *GAD67Cre* mouse pups we injected either a mix of 1.3μL of AAV2.1-hSyn-GCAMP6s.WPRE.SV40 with 0.7μL of AAV9-FLEX-CAG-tdTomato or 2μL of AAV9-hSyn-FLEX-axon-GCaMP6s.

#### Intra-hippocampal injection

When hippocampal viral injections were performed at P0 (AAV1-hSyn-FLEX-nGToG-WPRE3), mouse pups were anesthetized by inducing hypothermia on ice and maintained on a dry ice-cooled stereotaxic adaptor (Stoelting, # 51615) with a digital display console (Kopf, Model 940). Dorsal hippocampus was targeted by empirically determined coordinates, based on the Atlas of the Developing Mouse Brain (Paxinos et al., 2020), using transverse sinus and superior sagittal sinus as reference: 0.8 mm anterior from the sinus intersection; 1.5 mm lateral from the sagittal sinus; 1.1 mm depth from the skull surface. Under aseptic conditions, an incision was made in the skin, the skull was exposed and gently drilled (Ball Mill, Carbide, #¼ .019” -0.500 mm diameter-, CircuitMedic). Ten nL of undiluted viral solution were injected using an oil-based pressure injection system (Nanoject III, Drummond Scientific, rate of 5 nL/min). The tip of the pipette was broken to achieve an opening with an internal diameter of 30-40 μm. When hippocampal viral injections were performed at P5 or P9 (SAD-B19-RVdG-mCherry), pups were anesthetized using 3 % isoflurane in a mix of 90 % O_2_-10 % air and maintained during the whole surgery (∼0:30 h) between 1 % and 2.5 % isoflurane. Body temperature was monitored and maintained at 36°C. Analgesia was controlled using Buprenorphine (0.05 mg/ kg). Under aseptic conditions, an incision was made in the skin, the skull was exposed, antero-posterior and medio-lateral coordinates of the dorsal hippocampus were estimated by eye looking at the skull sutures. The skull was gently drilled and 10 nL of a viral solution were injected (Nanoject III, Drummond Scientific, rate of 5 nL/min) at a depth of 1.25 mm below the dura.

#### Window implant surgery

The surgery to implant a 3-mm-large cranial window above corpus callosum was adapted from previous methods (Villette et al., 2015). Anesthesia was induced using 3 % isoflurane in a mix of 90 % O_2_-10 % air and maintained during the whole surgery (∼1:30 h) between 1 % and 2.5 % isoflurane. Body temperature was controlled and maintained at 36°C. Analgesia was controlled using Buprenorphine (0.05 mg/ kg). Coordinates of the window implant were visually estimated. Then a small custom-made headplate was affixed with cyanoacrylate and dental acrylic cement. The skull was removed and the cortex was gently aspirated until the appearance of the external capsule/alveus. At the end of the cortectomy, we sealed a 3 mm glass window diameter circular cover glass (#1 thickness, Warner Instrument) attached to a 3 mm diameter large and 1.2 mm height cannula (Microgroup INC) with Kwik-Sil adhesive (WPI) and fixed the edge of the glass with cyanoacrylate. We let the animal recover on a heated pad for at least one hour before the imaging experiment.

#### Imaging

Two-photon calcium imaging experiments were performed on the day of the window implant using a single beam multiphoton pulsed laser scanning system coupled to a microscope (TriM Scope II, LaVision Biotech). The Ti:sapphire excitation laser (Chameleon Ultra II, Coherent) was operated at 920 nm. GCaMP fluorescence was isolated using a bandpass filter (510/25). Images were acquired through a GaAsP PMT (H7422-40, Hamamatsu) using a 16X immersion objective (NIKON, NA 0.8). Using Imspector software (LaVision Biotech), the fluorescence activity from a 400 μm x 400 μm field of view was acquired at approximately 9 Hz with a 1.85 μs dwell time per pixel (2 μm/pixel). Imaging fields were selected to sample the dorsal CA1 area and maximize the number of imaged neurons in the *stratum pyramidale*. Piezo signal, camera exposure time, and image triggers were synchronously acquired and digitized using a 1440A Digidata (Axon instrument, 50 kHz sampling) and the Axoscope 10 software (Axon instrument). During the imaging session, body temperature is continuously controlled.

#### Behavioral recordings

Simultaneously with imaging experiments, mouse motor behavior was monitored. In a first group of animals, motor behavior was monitored using two or three piezos attached to the paws of the animal. The signal from the piezo was acquired and digitized using a 1440A Digidata and the Axoscope 10 software. In a second group of animals, pups were placed and secured on an elevated platform (with the limbs hanging down on each side without touching the ground nor the support, as described here (Blumberg et al., 2015)). Motor behavior was monitored using two infrared cameras (Basler, acA1920-155um) positioned on each side of the animal. For each camera, a square signal corresponding to the exposure time of each frame from the camera was acquired and digitized using a 1440A Digidata and the Axoscope 10 software.

#### Recording of electromyogram activity in neonatal mice

The vigilance state of neonatal mice was assessed through analysis of electromyogram (EMG) signals obtained from a single insulated Tungsten wire (A-M Systems 795500) implanted in the nuchal muscle. A stainless-steel wire (A-M Systems 786000) wire inserted on the skull surface above the cerebellum and secured in place with dental cement served as the reference electrode. Signals from the electrodes were first passed through a headstage pre-amplifier before being digitized at 16000 Hz (Digital Lynx SX, Neuralynx (the pre-amplifier and digitizer were both from Neuralynx, as was the acquisition software (Cheetah))) and saved to a hard disk. TTL signals from the imaging and camera acquisition systems were simultaneously recorded as well to enable precise synchronization of EMG recordings with the camera and imaging data.

#### In vivo extracellular electrophysiological recordings

Multisite probes (16 channels silicon probes with 50 um separation distance, NeuroNexus, USA) were used to record electrophysiological activity below the window implant and in the intact hippocampus. To do so, we positioned the mouse pup (that had previously undergone a window implant) on the experimental setup. To head fix the animal the skull surface was covered with a layer of dental acrylic except the area above the intact hippocampus. In the intact (contralateral) hippocampus the electrodes were positioned using the stereotaxic coordinates of approximately 1.5 mm anterior to lambda and 1.5 mm lateral from the midline. Hippocampus under the window was recorded through the hole drilled in the window implant. Both multisite silicon probes were positioned at the depth to record *strata oriens* (SO), *pyramidale* (SP), *radiatum* (SR) and *lacunosum moleculare* (SLM). After the positioning of the electrodes, the animal was left in the setup for one hour to recover followed by 2 hours recordings of the neuronal activity in both hippocampi simultaneously.

#### Histological processing

Pups were deeply anaesthetized with a mix of Domitor and Zoletil (0.9 and 60 mg/kg, respectively), then transcardially perfused with 4 % PFA in 0.1M PBS (PBS tablets, 18912-014, Life technologies). For perisomatic innervation analysis, brains were post-fixed overnight at 4°C in 4 % PFA in 0.1M PBS, washed in PBS, cryo-protected in 30 % sucrose in PBS, before liquid nitrogen freezing. Brains were then sectioned using a cryostat (CM 3050S, Leica) into 50 μm thick slices collected on slides. Sections were stored at -20°C until further usage. For tracing experiments, brains from *GAD67Cre* and *Emx1Cre* pups, were post-fixed overnight at 4 °C in 4 % PFA in 0.1M PBS, washed in PBS and sectioned using a vibratome (VT 1200 s, Leica) into sagittal 70–80 μm thick slices.

Sections were stored in 0.1M PBS containing 0.05 % sodium azide until further usage. Immunocytochemistry was processed as described previously (Bocchio et al., 2020). Briefly, sections were blocked with PBS-Triton (PBST) 0.3 % and 10 % normal donkey serum (NDS), and incubated with a mix of up to three primary antibodies simultaneously diluted in PBST with 1 % NDS overnight at room temperature with the following primary antibodies: rabbit anti-dsRed (1:1000; Clontech, AB_10013483), chicken anti-GFP (1:1000, Aves Labs, GFP-1020, AB_10000240), mouse anti-synaptotagmin-2 (1:100; Developmental Studies Hybridoma Bank, AB_2315626). After several washes, according to the mixture of primary antibodies, the following secondary antibodies were used: donkey anti-chicken Alexa 488 (1:500, SA1-72000), donkey anti-rabbit Alexa 555 (1:500, ThermoFisher, A31570), donkey anti-mouse Alexa 488 (1:500, ThermoFisher, A21202), donkey anti-mouse Alexa 647 (1:500, ThermoFisher, A31571). After Hoechst counterstaining, slices were mounted in Fluoromount. Epifluorescence images were obtained with a Zeiss AxioImager Z2 microscope coupled to a camera (Zeiss AxioCam MR3) with an HBO lamp associated with 470/40, 525/50, 545/25, and 605/70 filter cubes. Confocal images were acquired with a Zeiss LSM-800 system equipped with a tunable laser providing excitation range from 405 to 670 nm. For quantifying synaptotagmin-2, 11μm thick stacks were taken (z=1μm, pixel size=0,156μm) with the confocal microscope using a plan-Achromat 40x/1,4 Oil DIC objective.

### DATA PREPROCESSING

#### Motion correction

Image series were motion corrected either by finding the center of mass of the correlations across frames relative to a set of reference frames (Miri et al., 2011) or using the NoRMCorre algorithm available in the CaImAn toolbox (Pnevmatikakis and Giovannucci, 2017) or both.

#### Cell segmentation

Cell segmentation was achieved using Suite2p (Pachitariu et al., 2017). Neurons with pixel masks including processes (often the case for interneurons located in the stratum oriens) were replaced by soma ROI manually drawn in ImageJ and matched onto Suite2p contours map using CICADA (**C**alcium **I**maging **C**omplete **A**utomated **D**ata **A**nalysis, source code available on Cossart lab GitLab group ID: 5948056). In experiments performed on *GAD67Cre* animals, tdTomato labelled interneurons were manually selected in ImageJ and either matched onto Suite2p contours map or added to the mask list using CICADA.

#### Axon segmentation

Axon segmentation was performed using pyAMNESIA (a **Py**thon pipeline for analysing the **A**ctivity and **M**orphology of **NE**urons using **S**keletonization and other **I**mage **A**nalysis techniques, source code available on Cossart lab GitLab group ID: 5948056). pyAMNESIA proposes a novel image processing method based on 3 consecutive steps: 1) extracting the axonal morphology of the image (skeletonization), 2) discarding the detected morphological entities that are not functional ones (branch validation), and 3) grouping together branches with highly correlated activity (branch clustering). To extract the skeleton, we first perform 3D gaussian smoothing and averaging of the recording, producing an image that summarizes it; on this image are then successively applied a histogram equalization, a gaussian smoothing, an adaptive thresholding, and finally a Lee skeletonization (Suen et al., 1994), allowing for the extraction of the skeleton mask and the morphological branches. To ensure the functional unity of the segmented branches, we only kept those that illuminated uniformly, where uniformity was quantified by the skewness of the pixel distribution of the branch during a calcium transient (branch validation). To cluster the valid branches based on their activity, we first extracted their average trace -- being the average image intensity along the branch for each frame -- and then clustered the branches traces using t-SNE and HDBSCAN algorithms with Spearman correlation metric (branch clustering).

#### Cell type prediction

Cell type prediction was done using the DeepCINAC cell type classifier (Denis et al., 2020). Briefly, a neuronal network composed of a convolutional neural network (CNN) and Long Short-Term Memory (LSTM) was trained using labeled interneurons, pyramidal cells and noisy cells to predict the cell type using 100 frames long movie patches centered on the cell of interest. Each cell was classified as interneuron, pyramidal cell or noise. Cells classified as ‘noisy cells’ were removed from further analysis. ‘Labelled interneurons’ were first kept in a separate cell type category and added to the interneurons list.

#### Activity inference

Activity inference was done using DeepCINAC classifiers (Denis et al., 2020). Briefly, a classifier composed of CNN and LSTM was trained using manually-labelled movie patches to predict neuronal activation based on movie visual inspection. Depending on the inferred cell type, activity inference was done using either a general classifier or an interneuron-specific classifier. Activity inference resulted in a (cells x frames) matrix giving the probability for a cell to be active at any single frame. We used a 0.5 threshold in this probability matrix to obtain a binary activity matrix considering a neuron as active from the onset to the peak of a calcium transient.

#### Behavior

Piezo signals were manually analyzed in a custom-made graphical user interface (Python Tkinter) to label the onset and offset of ‘twitches’, ‘complex movements’ and ‘unclassified movements’. Twitches were defined as brief movements (a few hundred milliseconds-long) occurring within periods of rest and detected as rapid deflections of the piezo signal. ‘Complex movements’ were defined as periods of movement lasting at least 2 seconds; ‘unclassified movements’ were short movements that differed from twitches and complex movements. Analysis of video tracking was done using CICADA and behavior was manually annotated in the BADASS (Behavioral Analysis Data And Some Surprises) GUI.

#### Neurodata without border (NWB: N) embedding

For each imaging session, imaging data, behavioral data, cell contours, cell type prediction, calcium traces and neuronal activity inference were combined into a single NWB: N file (following the guidelines provided on https://www.nwb.org/). Our NWB: N data set is accessible on DANDI archive (https://gui.dandiarchive.org/#/) - ref DANDI:000219.

### MODELING

#### Network implementation

We constructed a simple rate model and subsequently a more realistic spiking network in order to test our hypothesis that an increase in perisomatic inhibition could explain the switch in network dynamics between the first and second postnatal week. Both models consisted of one excitatory and one inhibitory population with recurrent interactions (**Figure 5A**, supplementary methods). The development of perisomatic innervation was simulated by increasing the strength from inhibitory to excitatory cells (J_EI_). The external input to the model was composed of a constant and a white noise term. To estimate the responses to twitch-like inputs, an additional feedforward input composed of short pulses was fed to the network. In the rate model, the rates represented the population averaged activities. The spiking network was constructed with Leaky Integrate and Fire (LIF) neurons. The network connectivity was sparse and each neuron received inputs from randomly selected neurons. Presynaptic spikes resulted in exponentially decaying postsynaptic currents. All codes used for the modelling are available at https://gitlab.com/rouault-team-public/somatic-inhibition/

### DATA ANALYSIS

#### Sample-size estimation

This study being mainly exploratory in the sense that the evolution of population activity in the CA1 region of the hippocampus using large scale imaging has not been described before, we have not been able to use explicit power calculation based on an expected size effect.

#### Histological quantifications

Confocal images of synaptotagmin-2 immunostaining were analyzed using RINGO (RINGs Observation), a custom-made macro in Fiji. We first performed a max-intensity projection of the Z-stack images of the top 6 μm from the slice surface, then images were cropped to restrict the analysis to the pyramidal cells layer. Obtained images were denoised using Fiji “remove background” option and then by subtracting the mean intensity of the pixels within a manually drawn ROI in the background area (typically the cell body of a pyramidal neuron). Denoised images were then binarized using a max-entropy thresholding (Fiji option). Finally, particles with size between 0.4 μm^2^ and 4 μm^2^ were automatically detected using the Fiji “Analyse particle” option. We then computed the proportion of the pyramidal cell layer (i.e., surface of the cropped region) covered by positive synaptotagmin-2 labelling.

#### In vivo electrophysiology

The neuronal activity recorded from both hippocampi in vivo using a 64-channel amplifier (DIPSI, France) was analyzed post-hoc. Firstly, data was downsampled to 1 kHz to save disc space. The local field potential (LFP) was band-passed (2-100 Hz) using the wavelet filter (Morlet, mother wavelet of order 6) and the common reference was subtracted to exclude the bias produced by volume conducted fluctuations of LFP. Sharp Wave events (SWs) were detected using a threshold approach. Firstly, LFP was bandpassed (2-45 Hz) and the difference between lfp recorded in the strati oriens, pyramidale and radiatum was calculated. Events were considered as SWs if: i) LFP reversion was observed in the stratum pyramidale, ii) their peak amplitude in the resulting trace exceeded the threshold of 4 standard deviations calculated over the entire trace (the threshold corresponds to p values below 0.01). The occurrence rate of SW was calculated over the entire recording and normalized to 1 min. SW co-occurrence was also calculated by cross correlating the SW timestamps from ipsilateral and contralateral hippocampi using a bin size of 10 milliseconds. Spectral analysis was carried out using the Chronux toolbox (Chronux, toolbox, 2010). Spectral power was estimated using direct multi-taper estimators (3 time-bandwidth product and 5 tapers).

#### Statistics for in vivo electrophysiology

Group comparisons were done using non parametric Wilcoxon rank sum test for equal medians, p-value of 0.05 was considered significant. Variability of the estimates was visualized as shaded bands of standard deviation computed using jackknife.

#### Vigilance state determination in neonatal mice

All analysis of EMG data was completed using custom scripts in MATLAB. For each experiment, the raw EMG data was first downsampled to 1000 Hz and subsequently high-pass filtered at 300 Hz and rectified. The processed data was then plotted to allow for manual inspection. Consistent with prior reports (Mohns and Blumberg, 2010), the data was primarily composed of alternating periods of high EMG tone (referred to as wakefulness) associated with ‘complex’ movements as well as periods of low EMG tone associated with a general behavioral quiescence and the presence of periodic brief myoclonic twitches (referred to as ‘active sleep’ due to the frequent observation of muscle twitches (Mohns and Blumberg, 2010). For vigilance state determination, we therefore utilized a protocol similar to that described previously (Rio-Bermudez et al., 2015). For both the ‘high’ and ‘low’ EMG tone conditions, 5 periods, each 1 s in duration, were first sampled from locations spread out over the entire recording length. Data from the samples were then pooled for each condition and the average value of the rectified signal was determined. Next, the midpoint between the average rectified signal values calculated for the ‘high’ and ‘low’ EMG tone conditions was determined for subsequent use as a threshold to separate periods of non-wakefulness (below the midpoint threshold value) from periods of wakefulness (above the midpoint threshold value), while the quarter point between these two values was calculated to further separate periods of non-wakefulness into active sleep (below the quarter point threshold value) or a sleep-wake transitory state (above the quarter point threshold value but below the midpoint threshold value). Once these thresholds were determined, the entire length of data was divided into 1 s non-overlapping bins and the average filtered rectified EMG signal was determined for each. A hypnogram was then created by automatically applying the threshold-derived criteria to the binned averaged data. Data bins scored as being active sleep were further analyzed to determine the presence of muscle twitches; this was accomplished by automatically identifying those bins whose filtered rectified average value exceeded 5 x the mean value determined from the low EMG tone representative samples. As a final step, the hypnogram and filtered rectified EMG signal data were plotted and manually inspected to ensure the accuracy of results. The hypnogram was then incorporated in the final NWB: N file to serve in the definition of the epochs of wakefulness and active sleep.

#### Analysis of calcium imaging data

Analysis was performed using CICADA, a custom-made python toolbox allowing for the automatic analysis of calcium imaging data in the NWB format (Cossart lab Gitlab, group ID 5948056, Project ID: 14048984). CICADA can be installed following the installation guidelines presented at https://gitlab.com/cossartlab/cicada. Each figure panel resulting from an analysis performed in CICADA is associated with a configuration file provided as supplemental data. These configuration files can be loaded in CICADA with the option ‘Load a set of parameters’ allowing for the replication of the analysis.

#### Calcium transient frequency analysis

Analysis launched from CICADA ‘Transient’s frequency’ analysis. The transient frequency for each cell was computed using the count of calcium transient onsets divided by the duration of the recording and was then averaged across all cells imaged in one given mouse pup across one or more imaging sessions.

#### Synchronous Calcium Event (SCE) detection

Analysis launched from CICADA ‘SCE description’ analysis. SCEs were defined as the imaging frames within which the number of co-active cells exceeded the chance level as estimated using a reshuffling method. Briefly, an independent circular shift was applied to each cell to obtain 300 surrogate raster plots. We computed the 99^th^ percentile of the distribution of the number of co-active cells from these surrogates and used this value as a threshold to define the minimal number of co-active cells in a SCE. Peak of synchrony above this threshold separated by at least 5 imaging frames (500 ms) were defined as SCE frames. To compute the percentage of transients within SCEs we counted, for each cell, the number of its calcium transients (from onset to peak) crossing SCE frames and divided it by its total number of calcium transients. We averaged the obtained values over all the cells imaged per animal.

#### Peri-Movement-Time-Histograms (PMTH)

Analysis launched from CICADA ‘Population level PSTH’ analysis. A 20-second-long time-window centered on movement onset was used. For each movement within an imaging session, the number of cells activated or the sum of all cell’s fluorescence was calculated for each time bin in that 20-second-long window. We obtain as many values as movements per time bin; for each individual imaging session the 25th, the median and 75th percentiles of the distributions of these values per time bin are computed and divided by the number of imaged cells. To display the percentage of active cells at a given time bin, these values were multiplied by 100. To combine imaging sessions in an age group (*i*.*e*., P5, 6, 7, 8, 9, 10, 11, 12) all the median PMTHs from individual imaging sessions belonging to the given group were stacked and we represented at each time bin the 25th percentile, the median and the 75th percentile value of these median PMTHs. To evaluate chance level around movement onsets, 500 surrogate raster plots per imaging session were computed, and the above procedure was used to obtain chance level in each imaging session and then grouped. PMTH obtained from fluorescence signals were built from DF/F calcium traces.

#### Movement-related inhibition

Analysis launched from CICADA ‘Activity ratio around epochs’ analysis. A 4 seconds long window centered on the onset of movements was used. The total number of cells activated during this time period was calculated. If less than 40 % of these cells were activated within 2 seconds following movement onset, the movement was classified as an ‘inhibiting’ movement. This procedure was applied to all detected movements to obtain for each mouse pup the proportion of ‘inhibiting’ movements.

#### Movement- and immobility-associated cells

Analysis launched from CICADA ‘Epoch associated cells’ analysis. The number of transients per cell occurring during movement or immobility was calculated. These transient onsets were then circularly shifted 100 times and the same calculation was performed on each roll. We used the 99^th^ percentile of this distribution as a threshold above which the cell was considered as associated with movement or immobility. Finally, the proportion of cells associated with rest or immobility was calculated for each imaged mouse.

### Statistics

Statistical tests were performed using GraphPad (Prism).

## Supplementary methods

Perisomatic inhibition onto pyramidal neurons in CA1 is established during early stages of development. Firstly, we investigated the effect of perisomatic inhibition on response to twitch-like inputs in a rate model. It allowed us to analytically characterise and compare the changes in the dynamics of the model under the conditions of weak and strong inhibition. Secondly, we show through numerical simulations that similar changes can be observed in a more realistic spiking network model. We show that increasing the strength of perisomatic inhibition in our models can account for the experimentally observed decrease of responses to twitch like feedforward inputs.

### Rate model

The rate model consists two populations (Excitatory and Inhibitory) with interaction strengths *J*_*ab*_’ (*a, b* ∈ {*E, I*}, *J*_*ab*_ > 0). They receive feedforward input of strengths *J*_*a*0_> 0, from an external excitatory population with average firing rate *r*_0_. The rates (*r*_*E*_, *r*_*I*_) represent the population averaged activities. The time scale of their evolution is determined by τ_*E*_ and τ_*I*_ and follow,

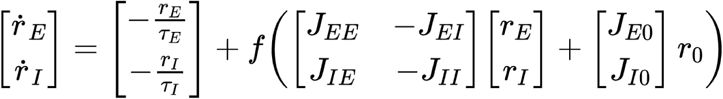

Where, *f*(*x*) is the neuronal transfer function. We choose it to be threshold linear. i.e., *f*(*x*) = [*x*] _+_. For positive *x*, the dynamics in matrix notation can be written as,

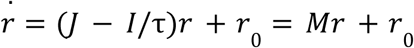

The fixed point of this system is given by,

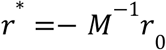

Where,

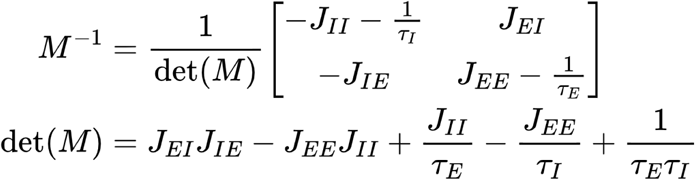

### Linear stability

The fixed point *r*^*^ is stable to small perturbations if the real part of all the eigenvalues of the Jacobian matrix *M*(*r*^*^) are negative. The eigenvalues λ_±_ can be expressed as,

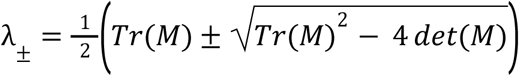

where,

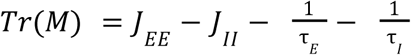

Equivalently, the system is always stable if *det*(*M*) > 0 and *Tr*(*M*) < 0.

Requiring, 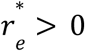 and 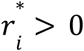 gives the conditions,

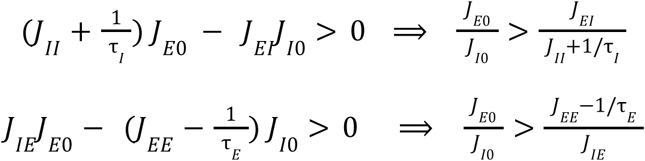

*det*(*M*) > 0 gives,

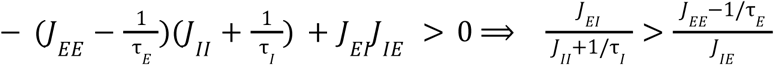

Combining the inequalities above gives the constraints for stable non-zero rates,

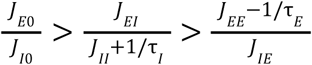

When the solutions are stable, small perturbations will decay to zero. Twitches can be considered as perturbations around the fixed point. The transient response to such short impulses can be expressed as *r*(*t*) = *C*_1_ *exp*(λ_+_) + *C*_2_ *exp*(λ_2_).

For a system with external white noise, ξ(*t*) as input we have,

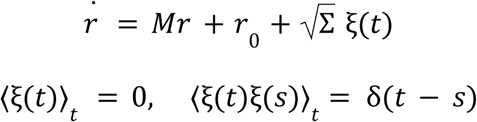

we set the off diagonal elements to zero, i.e, Σ_*ij*_ = 0 if *i* ≠ *j*.

Let δ*r*(*t*) = *r*(*t*) − *r*^*^, so the linearized system is,

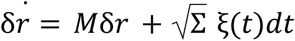

This can be seen as a 2D Ornstein - Ulenbeck process defined as: *dx*(*t*) =− *A x*(*t*) *dt* + *B dW*(*t*), which is well documented (Gardiner, 1985, p 109-111) and we can immediately write down the expression for the covariance matrix *C*(τ). With (*A* =− *M*) we have,

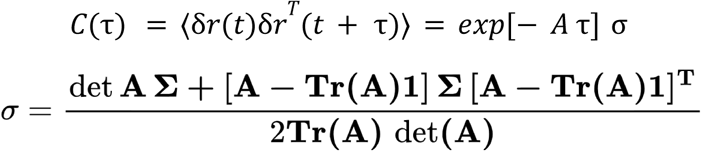

### Spiking model: LIF network

Excitatory and inhibitory neurons are modelled as Leaky Integrate and Fire (LIF) neurons. The LIF network consists *N*_*E*_ excitatory neurons and *N*_*I*_ inhibitory neurons with exponentially decaying postsynaptic currents. Each neuron receives exactly *K*_*E*_ excitatory and *K*_*I*_ inhibitory inputs from randomly selected neurons in the network. And we assume that the network is sparse i.e. *K* << *N*.

The evolution of sub-threshold membrane voltage 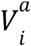 of neuron *i* in population *a* ∈ {*E, I*}, is given by,

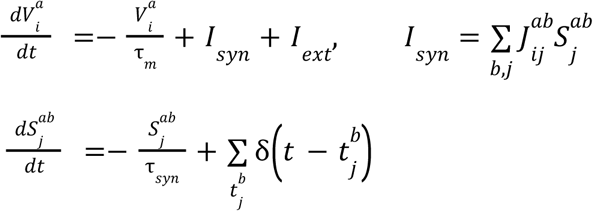

When the membrane voltage reaches the threshold, *V*_*thresold*_, it is reset to *V*_*reset*_. τ_*syn*_ is the synaptic time constant, 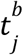 is the spike time of neuron (*i, b*). The coupling strengths are 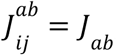, if there is a connection from neuron (*j, b*) to (*i, a*) and zero otherwise. The contribution of external inputs is represented by *I*_*ext*_. *I*_*syn*_ represents the total synaptic currents due to spikes. Spikes are modelled as delta functions.

If a neuron (*j, b*) emits a spike at time 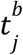 and projects to a post-synaptic neuron (*i, a*), this will result in a change of the membrane voltage 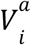 of the post-synaptic neuron by an amount *J*_*ab*_ in a time τ_*syn*_. The membrane voltage decays exponentially to its resting potential in a time τ_*m*_.

### Simulations and data analysis

The network simulations were conducted using custom code written in Python and C++ and all the analysis was done in Python. We use the forward Euler method to solve the set of coupled ode’s with a time step of 0.1ms.

### Covariance

Given a stationary stochastic process *X*_*t*_ with mean μ_*X*_ = *E* [*X*_*t*_], the autocovariance is given by,

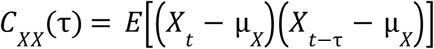

The covariance of *X*_*t*_ with another process *Y*_*t*_ is defined as,

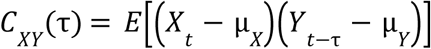

## ACKNOWLEDGMENTS

This work was supported by the European Research Council under the European Union’s, Horizon 2020 research and innovation program Grant#646925, the Fondation Bettencourt Schueller, and a NWB seed grant # R20046AA. The project leading to this publication has received funding from the « Investissements d’Avenir » French Government program managed by the French National Research Agency (ANR-16-CONV-0001) and from Excellence Initiative of Aix-Marseille University - A*MIDEX. R.F.D. was funded by the “Ministere de l’Enseignement Supérieur, de la Recherche et de l’Innovation” and by the Fondation pour la Recherche Médicale Grant FDT202106012824. E.L. was funded by the “Ministere de l’Enseignement Supérieur, de la Recherche et de l’Innovation” and by A*Midex foundation and the French National Research Agency funded by the French Government « Investissements d’Avenir » program (NeuroSchool, nEURo*AMU, ANR-17-EURE-0029 grant). J.D. was supported by the Fondation pour la Recherche Médicale Grant FDM20170638339. M.P was supported by the Fondation pour la Recherche Médicale Grant ARF20160936186. T. D and T. S. were funded by « Investissements d’Avenir » French Government program managed by the French National Research Agency (ANR-16-CONV-0001) and from Excellence Initiative of Aix-Marseille University - A*MIDEX. We would like to thank Dr. Pierre-Pascal Lenck-Santini for providing valuable feedback on our research project. We also would like to thank Marion Sicre for her help with *GAD67Cre* experiments. We are grateful to Pr. Hannah Monyer for providing the *GAD67Cre* mouse lines. We thank the Centre de Calcul Intensif d’Aix-Marseille for granting access to its high-performance computing resources. The rabies virus was a gift from Conzelman laboratory.

## AUTHORS CONTRIBUTION

RFD, MAP, and R.C designed research. RFD and EL performed stereotaxic injections in viral injections, RFD performed the surgery for window implant, RFD and MAP performed two photon calcium imaging experiments. CL, MK, AB, RFD and MAP performed histology experiments. MM, DS, RFD and MAP performed electrophysiological recordings, MM and DS analyzed these recordings. RFD and RB performed electromyogram recordings coupled with in vivo imaging and RB analyzed the EMG data. RFD, JD, MAP, and R.C designed the analysis pipeline. RFD and JD, wrote DeepCINAC, CICADA and RINGO algorithms. TS and TD wrote PyAmnesia. SRB and HR designed and simulated the network models. RFD, MAP and RC wrote the paper.

